# Post-loading ribonucleic acid into lipid nanoparticle carriers

**DOI:** 10.64898/2026.07.21.737377

**Authors:** Navid Bizmark, David F. Amelemah, Satya Nayagam, Sai Nikhil Subraveti, Bumjun Kim, Livia Salvati Manni, Kathleen Wood, Nageshwar Rao Yepuri, Michael Moir, Marina Cagnes, Nanzhi Zang, Gregory G. Warr, Robert K. Prud’homme

## Abstract

Lipid nanoparticles (LNPs) are effective carriers for messenger ribonucleic acid (mRNA) delivery in vaccines; however, their reliance on extreme cold-chain storage limits global manufacturing and distribution. Conventional LNPs are formed by rapidly mixing four lipids with mRNA through electrostatic interactions between cationic ionizable lipids and negatively charged nucleic acids, facilitating nucleation and precipitation of mRNA-loaded LNPs. However, this binding also accelerates mRNA degradation, requiring stringent cold storage which limits widespread vaccine deployment. To overcome this limitation, we introduce a *post-loading* strategy in which empty LNPs (eLNPs) are first fabricated and RNA is subsequently loaded at a later stage. Using scalable confined impinging jet (CIJ) mixers, we optimized pH, buffer composition, lipid concentration, and ethanol content to produce colloidally stable eLNPs. Controlled adjustment of ethanol content and pH enabled efficient incorporation of four distinct RNA payloads while maintaining loaded LNP diameters below 100 nm. Post-loaded LNPs demonstrated mRNA delivery efficiencies in HeLa cells comparable to those of conventionally co-precipitated LNPs. Consistent size distributions and zeta potentials further confirmed comparable surface properties. Structural characterization by x-ray and neutron scattering revealed similar internal architectures for post-loaded and co-precipitated LNPs without compromising RNA loading efficiency. Together, these results demonstrate equivalent cellular delivery performance between the two formulations. This post-loading approach enables decentralized assembly of mRNA LNPs at the point of administration, with both eLNPs and mRNA stored under mild refrigeration, thereby improving vaccine accessibility. Moreover, eLNPs function as modular laboratory reagents, facilitating the translation of mRNA research toward clinical applications.

## Introduction

The discovery of precise nucleoside modification in ribonucleic acid (RNA) has revolutionized RNA molecule synthesis, producing RNAs that have opened the field of messenger ribonucleic acid (mRNA) therapeutics.^1^ This breakthrough has paved the way for the design and development of RNA-based vaccines.^2^ These vaccines are engineered to safeguard RNA from enzymatic degradation during circulation in the body, ensuring maximal delivery to target cells while minimizing off-target effects.^3,4^ Among the various vectors explored,^4–7^ lipid-based nanoparticles have emerged as highly effective carriers for RNA delivery. The first RNA-based medication approved by the United States Food and Drug Administration (U.S. FDA)—marketed as Onpattro®—targets the treatment of peripheral nerve disease stemming from hereditary transthyretin-mediated amyloidosis (hATTR) in adult patients.^8,9^ In this technology, small interfering ribonucleic acid (siRNA) is encapsulated in lipid nanoparticles (LNPs) for systemic delivery to liver to reduce liver-produced TTR protein levels and subsequent amyloid deposits.^10^ Furthermore, LNPs played a crucial role in the development and deployment of mRNA-based vaccines for addressing the global health pandemic posed by coronavirus disease 2019 (COVID-19).^11–14^ In these vaccines, the therapeutic mRNA is encapsulated in LNPs for a systemic delivery through intravenous injection.

The production of RNA-loaded LNPs typically involves mixing an aqueous buffer stream containing the RNA cargo with an ethanol stream containing four lipids: (i) a cationic ionizable lipid, (ii) a zwitterionic lipid, (iii) cholesterol, and (iv) a pegylated lipid.^15^ The ionizable lipids utilized in these formulations typically possess a pKa ranging from 6 to 7,^16^ rendering them positively charged in acidic environments with a pH below their pKa. Under such conditions, the positively charged ionizable lipids form electrostatic complexes with the negatively charged RNAs, initiating the formation of nuclei during the mixing process. As mixing proceeds, these nuclei grow through the precipitation of zwitterionic lipid and cholesterol until reaching a critical point where the pegylated lipid assembles at the surface of the LNPs, halting further growth—a nucleation and growth mechanism.^17–19^ The mixing mode significantly influences the quality and efficacy of RNA-loaded LNPs.^20,21^ In instances of poor mixing, such as in bulk mixing, LNPs of approximately 150 nm diameter are produced with a low RNA encapsulation efficiency of around 60%.^22^ Conversely, when mixing is enhanced, particularly in microfluidic settings, much smaller particles of approximately 75 nm are generated with an encapsulation efficiency of approximately 90%, resulting in significantly improved in vitro and in vivo transfections. However, variations in microfluidic mixing parameters may yield LNPs with diverse physicochemical properties, affecting their delivery efficacy and organ tropism.^23^ Additionally, challenges such as fouling and limited throughput in microfluidic devices need to be addressed for scalable LNP production.^22,24^ Turbulent mixers have been demonstrated as a scalable route to LNP production^25,26^ and are the basis of all current commercial COVID vaccines.

The COVID-19 pandemic underscored the challenge of transitioning from laboratory-scale formulations to large-scale production capable of manufacturing over 3 billion vaccine doses. To meet this demand, a series of engineering designs and in-process controls were needed to ensure that the vaccines maintained the same quality observed during clinical trials.^27^ Pfizer’s manufacturing facilities achieved this goal by scaling the production of mRNA-loaded LNPs for COVID-19 vaccines, employing eight high-throughput turbulent mixers.^27^ The scalability of vaccine manufacturing was effectively addressed through these confined impinging jet (CIJ) mixers.^28–30^ However, mRNA vaccines also presented post-processing challenges due to their requirement of extreme cold chain storage and distribution. Both the Pfizer and Moderna vaccines were released with the requirement of handling temperatures of -70 °C and -40 °C, respectively.^27,31^ These freezing conditions are essential to suppress the oxidation and hydrolysis of mRNA, particularly when the mRNA phosphodiester bonds interact closely with the cationic ionizable lipid nitrogen groups co-assembled in LNPs.^32,33^ Interestingly, mRNA in a lyophilized state is stable at room temperature.^34^ Specifically, the degradation rate constant of lyophilized RNA stored in air- and moisture-tight containers at 25 °C is found to be 0.7-1.3 cuts per 1000 nucleotides per century, resulting in less than 10% RNA degradation over a 24-month period.^35^ Increased stability via lyophilization of LNPs has been an area of significant interest and activity. At the laboratory scale, lyophilized RNA-loaded LNPs with significant storage stability have been demonstrated.^36–39^

In contrast to the traditional approach of mRNA assembly in LNPs and then developing techniques to stabilize the formulation, we propose a two-component approach in which mRNA spontaneously assembles into stable “empty” lipid nanoparticles (eLNPs) at the time of administration. We denote the conventional mRNA LNP process as “co-loading” (**Figure 1A**), and our new approach as “post-loading” of eLNPs (**Figure 1B**). In this approach, lipid components are initially assembled into eLNPs in the absence of mRNA cargo, then frozen under mild conditions or lyophilized for long-term storage. Upon reaching the administration site, the eLNPs are either thawed or rehydrated in buffer and then mixed with rehydrated lyophilized mRNA of interest. This approach has several important advantages:

1. The stability of lipids in eLNP and of the lyophilized mRNA eliminates the requirement for extreme cold chain handling. This enables these mRNA LNP therapeutics to be used in low-resourced, global health applications, which is not possible with current mRNA LNP vaccines.
2. The rapid mutation of viruses such as coronavirus means that mRNA sequences need to be updated.^40^ Whereas co-loaded mRNA LNPs must be abandoned whenever new mRNA sequences are required, in the post-loading approach new lyophilized mRNA can be used to prepare mRNA LNPs without loss.
3. Post-loading enables confident translation from laboratory research to clinical applications. In many biologically oriented laboratories mRNA is combined with cationic reagents such as Lipofectamine® to assemble RNA-LNPs.^41,42^ While effective in laboratory settings, cytotoxic Lipofectamine® cannot be utilized in human clinical formulations. In contrast, eLNPs can be formulated with lipid components approved by the U.S. FDA in current LNP vaccines, allowing their use as a reagent to complex mRNA for rapid screening. Therefore, transfection pathways, cytotoxicity, and immune responses at the laboratory level are far more likely to translate to clinical use.

**Figure 1.**
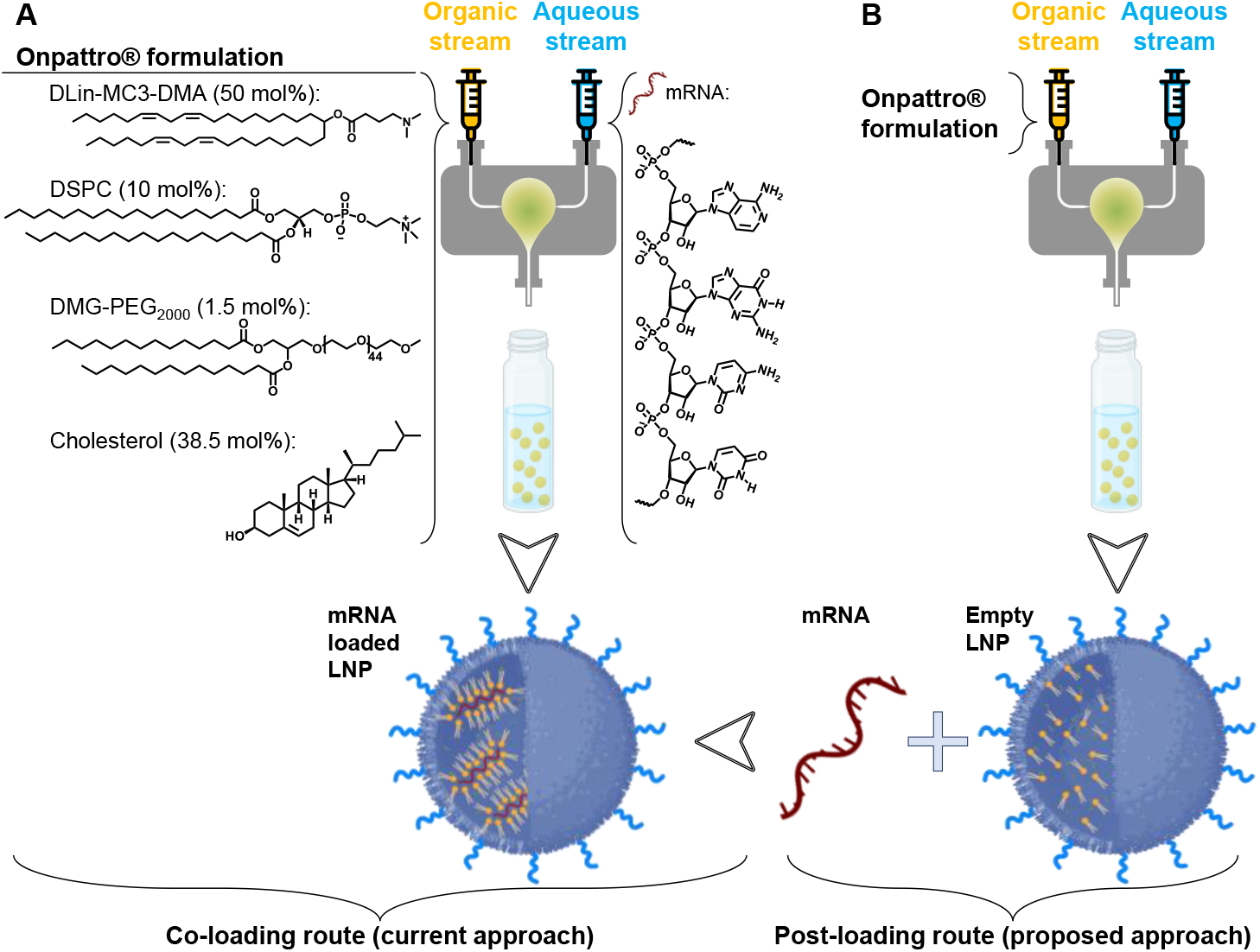
Schematic of lipid nanoparticle production via flash nanoprecipitation. **A**. The schematic of a confined impinging jet (CIJ) mixer where an ethanol solution of lipid components (organic stream), comprised of cationic ionizable lipid (DLin-MC3-DMA, in short MC3), zwitterionic phospholipid (DSPS), cholesterol, and PEGylated lipid (DMG-PEG_2k_) at molar ratios as noted (i.e., Onpattro® formulation) was quickly mixed with an aqueous stream containing nucleic acid in an appropriate buffer at a pH below the pK_a_ of MC3 (i.e., 6.44). After mixing, the mixture was collected and diluted in a quench bath to reach 5 vol.% ethanol. We call this route of production “co-loading” as lipids and nucleic acid are precipitated together, and LNPs are loaded with RNA at the time of formation. **B**. Lipid precipitation route as in A but in the absence of nucleic acid in the aqueous stream. After the formation of “empty” LNPs, RNA was added under controlled ethanol content and pH to develop a “post-loading” route of production.

The concept of post-loading appears to have first been presented in the patent by Moderna entitled “Methods of Preparing Lipid Nanoparticles”,^43^ and has been recently used in vaccine formulations.^44,45^ However, little of the physical chemistry and formulation background has been provided. In this contribution we first present how eLNPs can be produced using scalable CIJ mixers and discuss what factors impact their physicochemical properties and colloidal stability. In CIJ mixers, an organic solvent stream containing the lipids and an aqueous anti-solvent stream are rapidly mixed at a Reynolds number of ∼15,000.^28^ The mixing process is crucial for the successful production of eLNPs. As a model we use a formulation comprised of U.S. FDA-approved lipids of (6Z,9Z,28Z,31Z)-heptatriacont-6,9,28,31-tetraene-19-yl 4-(dimethylamino)butanoate (DLin-MC3-DMA, or in short MC3), 1,2-distearoyl-sn-glycero-3-phosphocholine (DSPC), cholesterol, and 1,2-dimyristoyl-rac-glycero-3-methoxypolyethylene glycol-2000 (DMG-PEG_2k_) (Figure 1). This is generically called the Onpattro® formulation. MC3 is a cationic ionizable lipid (pKa 6.4), which is >90% protonated at or below pH 5.5 but only ∼25% charged at pH 7.^16,46–48^ In the current co-loading technology (Figure 1A), MC3-mRNA electrostatic complexation initiates the nucleation of LNP formation during the rapid mixing of four lipids against an aqueous solution of mRNA buffered at pH 4-5.5 resulting in the entrapment of ∼24% water within LNPs.^49^ In the alternative technology (**Figure 1B**), we make eLNPs first and then post-load them with RNAs. We discuss how the two factors of (i) electrostatic attraction between eLNPs and RNAs and (ii) lipid mobility can be tuned to achieve at 80%-95% encapsulation efficiency for RNA of various payloads while maintaining the size of post-loaded LNPs below 100 nm. Such post-loaded LNPs demonstrate similar protein expressions when compared to co-loaded LNPs in HeLa cell cultures. Small-angle x-ray scattering (SAXS) and small-angle neutron scattering (SANS) experiments together with cryo-electron microscopy (cryo-EM) are presented to compare the structures of the eLNPs, as well as co-loaded and post-loaded LNPs.

## Materials and Methods

### Materials

We purchased purified *Torulla Ambion*™ yeast RNA (<170 nt)^50^ from ThermoFisher Scientific, CleanCap® Firefly Luciferase messenger RNA (1929 nucleotides, fully substituted with 5-methoxyuridine) (FLuc mRNA) from TriLink Biotechnologies, dasher green fluorescent protein (GFP) mRNA (1050 nt) from Aldevron, and carrier RNA (Poly A) from QIAGEN. For lipid nanoparticle formulations, we purchased (6Z,9Z,28Z,31Z)-heptatriacont-6,9,28,31-tetraene-19-yl 4-(dimethylamino)butanoate (DLin-MC3-DMA, or in short MC3) from Nanosoft Polymers, 1,2-distearoyl-sn-glycero-3-phosphocholine (DSPC), 1,2-dimyristoyl-rac-glycero-3-methoxypolyethylene glycol-2000 (DMG-PEG_2k_), and cholesterol from Avanti Polar Lipids.

Deuterated lipids were synthesized as described in what follows. Perdeuterated 1,2-distearoyl-d_70_-*sn*-glycero-3-phosphocholine-d_13_ (DSPC-d_83_ average 96.5%D, see the Synthesis of Deuterated Materials section and Schematic S1 in the Supporting Information) was synthesized from stearic acid. Hydrothermal deuteration of the saturated fatty acid starting material afforded high deuterium incorporation (96%D) after two cycles.^51^ The deuterated fatty acid was coupled with 3-*O*-benzyl-*sn*-glycerol under Steglich esterification conditions. Removal of the benzyl ether protecting group, followed by phosphorylation and 2,4,6-triisopropylbenzenesulfonyl chloride-mediated coupling with choline-d_13_ tetraphenylborate furnished the desired deuterated phospholipid (see Figures S7 and S8 in the Supporting Information for the molecular analyses). 4-(Dimethylamino)-butanoic acid, (10Z,13Z)-1-(9Z,12Z)-9,12-octadecadien-1-yl-10,13-nonadecadien-1-yl ester-d_62_ (DLin-MC3-DMA-d_62_ average 96%D, see the Synthesis of Deuterated Materials section and Schematic S2 in the Supporting Information) with deuterium labelling in the linoleoyl-derived tails was synthesised from linoleic acid-d_31_,^52^ using modified procedures.^16^ Briefly, linoleic acid-d_31_ was converted to the corresponding alkyl bromide and treated with magnesium to afford the requisite Grignard reagent. Reaction with ethyl formate and coupling of the resulting secondary alcohol with 4-(dimethylamino)butanoic acid provided the tail deuterated ionisable lipid (see Figures S9-S11 in the Supporting Information for the molecular analyses). Uniformly deuterated cholesterol-d_45_ (average 79%D) was produced as previously reported (see Figures S12-S14 in the Supporting Information for the molecular analyses).^53^

For pH 7 buffers, 2-[4-(2-Hydroxyethyl)piperazin-1-yl]ethanesulfonic acid (HEPES free acid, IBI Scientific) was used. For acidic buffers (pH 4, 5, and 5.5), acetic acid (Sigma-Aldrich) and sodium acetate (Aldrich) at the desired ratios were used with a complementary amount of 1 M sodium hydroxide (Fisher Chemical) to adjust the final pH. All the aqueous and organic solutions were prepared in nuclease-free water (Qiagen) and ethanol (ThermoFisher Scientific), respectively. When needed, LNPs were made in D_2_O buffer.

### Lipid nanoparticle (LNP) formulations

We utilized the CIJ technique,^17,28^ to fabricate both eLNPs and co-loaded LNPs *via* flash nanoprecipitation (FNP). To produce co-loaded LNPs, we employed an Onpattro-based formulation^54^ consisting of MC3/DSPC/cholesterol/DMG-PEG_2k_ at a molar ratio of 50/10/38.5/1.5 in ethanol, respectively. This ethanol stream was mixed against an equivalent volume of an aqueous buffer stream containing the desired RNA using a CIJ mixer (**Figure 1A**). The total lipid mass concentration was maintained at 12 mg/ml in the ethanol feed stream, with a molar ratio of ionizable lipid to RNA set at 6 for optimal in vivo efficacy.^55–57^ To produce eLNPs, we repeated the mixing procedure without the presence of RNA (**Figure 1B**). After mixing, the resulting co-loaded LNPs and eLNPs were collected in a buffer reservoir at a specified pH, ensuring that the final dispersion contained 5 vol.% ethanol. The ionic strength of the buffer was set at 20 mM in the jet stream and 10 mM in the reservoir, maintaining an effective ionic strength of 10 mM during mixing and collection. For post-loading LNPs, the desired RNA (in a pH 7 HEPES buffer) was initially added to the freshly prepared eLNP suspension at neutral pH then titrated to pH 5.5. Similar to co-loaded LNPs, the molar ratio of ionizable lipid to RNA in the post-loaded LNPs was kept at 6, unless mentioned otherwise. All subsequent analyses on post-loaded LNPs were performed under these conditions.

### Dynamic light scattering (DLS) and zeta sizing

The diameter and zeta potential of LNPs were assessed using DLS with an Anton Paar Litesizer 500 instrument. This instrument was equipped with a single-frequency laser diode generating laser light at a wavelength of 658 nm with a power output of 40 mW. By collecting backscattered light at an angle of 175°, we determined the size of LNPs utilizing the Stokes-Einstein model and the first cumulant of the series expansion of the light scattering correlation function. Zeta potential measurements of LNPs were conducted using folded capillary cells (Omega cuvettes) with the same equipment. Both size and zeta potential measurements were conducted independently at least three replicates, and the LNP suspensions were utilized as prepared in FNP without any further dilution.

### Ribogreen® assay for encapsulation efficiency (EE)

EE of RNA in LNPs was assessed using the Ribogreen® assay (Quant-iT Ribogreen® RNA Reagent, ThermoFisher Scientific).^58^ Following the manufacturer’s protocol, the Ribogreen® dye was diluted 200-fold in a Tris-EDTA (TE) buffer of pH 7.5, optimized for RNA concentration detection within the range of 0.02 to 1.0 µg/ml. LNP suspensions were then diluted in TE buffer to achieve a total mass concentration of approximately 0.6 µg/ml for RNA, with ethanol content below 0.2%. Identical samples were prepared with the addition of 0.5 wt.% Triton X-100 (Sigma Aldrich), which effectively dissociates the LNPs and releases RNA. This facilitated the distinction between “free” and total RNA concentrations in the absence and presence of Triton X-100, respectively, during the Ribogreen® assay. Calibration curves were constructed using RNA solutions provided in the assay kit, ranging from 0 to 1 µg/ml, and diluted in the same TE buffer at pH 7.5. Corresponding RNA solutions with 0.5 wt.% Triton X-100 were also prepared. Each solution, with or without Triton X-100, was dispensed in 100 µl volumes into separate cells of a 96-well, non-treated, flat-bottom, opaque plate for analysis using a SpectraMax i3x plate reader (Molecular Devices). Triplicate measurements were conducted for each LNP sample, while calibration curve RNA solutions underwent duplicate measurements. Fluorescence intensity was recorded at an excitation wavelength of 485 nm and an emission wavelength of 528 nm, with an illumination duration of approximately 10 seconds per reading. The EE was computed from 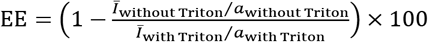, where *Ī* _with Triton_ and *Ī*_without Triton_ are the averaged fluorescence intensity of three replicates of samples without and with Triton X-100, respectively, and *a*_with Triton_ and *a*_without Triton_ denote the slope of linear fits to the two averaged calibration curves with and without Triton X-100, respectively.

### In vitro cell transfection assay

HeLa cells were purchased from American Type Culture Collection (ATCC) and cultured in complete Dulbecco’s modified Eagle’s medium (DMEM, HyClone, USA), supplemented with 10% fetal bovine serum (FBS) and 1% penicillin/streptomycin at 37 °C in 5% CO_2_. Cell passages were facilitated using 0.25% trypsin-EDTA (Gibco, USA). Approximately 20,000 cells were then seeded into 96-well plates, and Luc or GFP mRNA-loaded LNPs, whether co-loaded or post-loaded, were diluted with complete DMEM to a concentration ranging from 50 to 500 ng/ml mRNA within the plates. Dilutions were made with consideration of EE so that identical concentrations of encapsulated RNA were used. Each sample was prepared in triplicate or quadruplicate as indicated in the manuscript. As a positive control, cells were transfected with mRNA using Lipofectamine 3000 (Thermo Fisher Scientific, Waltham, MA, USA) following the manufacturer’s protocol. After 24 hours of incubation at 37 °C in a 5% CO_2_, treated cells were assessed for luciferase or green protein expression using Brightglo luciferase assay (Promega). To assess cell viability, resazurin sodium salt was used in place of Bright-Glo™. Cells were treated with a 50 μM concentration of resazurin sodium salt for a duration of 4 hours. The cell viability was then measured with a SpectraMax i3x Multi-Mode Microplate Reader, using excitation and emission wavelengths of 530 nm and 590 nm, respectively.

### Small angle x-ray scattering (SAXS)

Synchrotron SAXS experiments were conducted at the 16ID-LiX beamline of the National Synchrotron Light Source II at Brookhaven National Laboratory, USA. Standard flow-cell-based apparatus with quartz capillaries from Schott AG were employed for all measurements. The scattered data were collected using Pilatus3X 1M and Pilatus3X 900K upon exposure to 6-18 keV x-ray beam for an exposure time of 0.5 s. With a sample-to-detector distance of 3.562 m, the q range falls between of 0.005 A°^-1^ to 3.19 A°^-1^. The scattering vector, *q*, is defined as 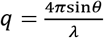, where 0 is the scattering angle and λ is the beam wavelength (i.e., 0.8189 A°). For each measurement, the sample was concentrated 2-3 times to achieve a concentration of 1 to 2 mg/ml for the total lipid concentration. To this aim, an appropriate volume of LNPs was centrifuged using 100 kDa MWCO Amicon® Ultra Centrifugal Filters (Millipore) at 1000 RCF. Both the filtrate and concentrated samples were utilized for SAXS measurements. Initially, empty cells and blank solvents (i.e., filtrate from the filtration step) were measured for proper background subtraction. Subsequently, 10 measurements were taken at various points across the sample. The resulting scattering profiles were obtained by averaging the repeated measurements and subtracting the contributions from the empty cell and blank solvent using the Python package py4xs in Jupyter Notebooks.

### Small angle neutron scattering (SANS)

SANS experiments were performed using the Quokka beamline at the ANSTO Australian Centre for Neutron Scattering, Australia.^59^ The data were collected in three different configurations to give the total scattering intensity over a scattering vector, *q*, range of 0.002 A°^-1^ to 0.6 A°^-1^. For the medium *q* and high *q* ranges, a neutron wavelength of 6 A° was used with sample to detector distances of 8 m (medium *q*) and 1.3 m (high *q*), and source to sample distances of 8 m and 4 m, respectively. To access lower *q* values, neutrons of 8 A° wavelength, sample to detector distance of 20 m and source to sample distance of 20 m were used. LNP samples were prepared as described earlier, but the hydrogenous MC3, DSPC, and cholesterol were replaced by deuterated MC3-d_62_, DSPC-d_83_, and cholesterol-d_45_, respectively, while maintaining their molar ratios similar to those of the hydrogenous components (

See the Synthesis of Deuterated Materials in the Supporting Information for more details). Two batches of deuterated LNPs were prepared, one in 100% H_2_O and the other in 100% D_2_O buffers. These batches were mixed at appropriate volumes to yield LNPs with different external contrasts (i.e., D_2_O/H_2_O ratios). The LNPs were then concentrated from 0.6 mg/ml to 3 mg/ml as described above. The measurements were done in Hellma quartz disk-shaped cuvettes of 1- and 2-mm path lengths depending on the H_2_O content of the sample. The samples were maintained at 28 °C throughout the measurements. The duration of the measurements was between 50 and 100 min per sample, depending on the count rate. The data from the different configurations were reduced for instrument effects using empty cell and blocked beam measurements, placed on an absolute scale using direct beam measurements with a calibrated attenuator and merged into a single file. Standard SANS reduction protocols were used in Igor Pro.^60^ The solvent was background subtracted using the correlation function within SasView. The analysis of the SANS data was done with SasView and is further described in the Supporting Information.

### Cryo-electron microscopy (cryo-EM)

QF-Cu300 R1.2/1.3 + 2 nm carbon grids were discharged for 7 s on CEMRC GloQube at 20 mA. 3 μL of sample was applied to each grid at the provided concentration. Vitrobot condition was set to temperature at 4 °C; humidity at 95 %; wait time for 45 s; drain time for 0.5 s; blot force for 13, and blot time for 3 or 5 s for all grids. Grids were then prepared on the CEMRC Vitrobot Mark IV set to 4 °C/95% humidity. All images collected on the Talos Arctica 200kV TEM equipped with a Gatan K3 camera and BioQuantum energy filter (20 eV) using SerialEM software. Images were collected from holes distributed across overview maps. Images were collected at higher magnification with a 1.063 A°/pixel and a defocus range of -2.0 to -3.0 micron, with a total 60 e-/A°^2^ electron dose.

## Results and Discussion

### Empty lipid nanoparticle (eLNP) formulation

The post-loading technology requires stable eLNPs as precursors (**Figure 1B**). During the rapid mixing process in CIJ mixers, the assembly of mRNA and ionizable lipids is influenced by the charge state of ionizable lipid, which depends on solution pH.^61^ mRNA LNP formulations are often characterized by their N:P ratio; i.e., the ratio of ionizable cationic nitrogen groups to anionic phosphate groups on the mRNA. However, pH relative to the pKa of the ionizable lipid determines the true charge ratio and hence controls the assembly. To explore these key factors, we formulated eLNPs under three mixing pH conditions: pH 4, pH 5, and pH 5.5. In all experiments, the mixture of aqueous and ethanol streams was quenched into a pH 7 buffer yielding a final 5 vol.% ethanol. An ionic strength of 10 mM was maintained during both the mixing and quenching steps (see Materials and Methods for details). The total lipid mass concentration in the ethanol stream was set at 12 mg/ml.

**Figure 2** illustrates the hydrodynamic diameter and zeta potential of the three formulations, together with their stability over time. In all cases, we achieved the production of eLNPs ranging from 50 to 65 nm with a polydispersity index (pdi) of less than 0.2 (**Figure 2A**). For all pH processing histories, eLNPs exhibited colloidal stability for at least three days when stored at room temperature (20-25 °C) or in a 4 °C refrigerator (see Figure S1 in the Supporting Information for the size distributions). At room temperature, the size increased by less than 6 nm after 3 days, whereas at 4 °C, it only increased by 3 nm during the same period. The final pH post-mixing and quenching was 7.0±0.1, at which approximately 25% of MC3 is positively charged,^48^ rendering the eLNPs slightly positively charged as evidenced by zeta potentials of approximately +10 mV (**Figure 2B**). Through a comparison of the computed Debye length and PEG thickness (See the Debye length and PEG brush thickness calculations in the Supporting Information), we infer that particle stability during processing is attributed to steric and electrostatic stabilization. The presence of PEG is essential to inhibit particle coalescence over time. While all eLNPs exhibited good colloidal stability at room temperature, those made at pH 4 with approximately 65 nm hydrodynamic diameter demonstrated superior stability. After three days, their diameter increased only by approximately 2 nm while maintaining pdi<0.15, likely due to the greater electrostatic repulsions between eLNPs at lower pH. However, eLNPs formulated at pH 5 possessed the smallest size of approximately 50 nm, and thus, used these to study the post-loading dynamics. Post-loading requires lowering the pH of the medium in which eLNPs are dispersed to at least one unit below the corresponding pKa of MC3.^48^ **Figure 2C** illustrates the size and zeta potential of eLNPs resulting from quenching at pH 7 after mixing at pH 5. When eLNPs were titrated down to pH 4.5 through the addition of an appropriate amount of 100 mM acetate buffer, the size of eLNPs remained unchanged during protonation (**Figure 2C**).

**Figure 2.**
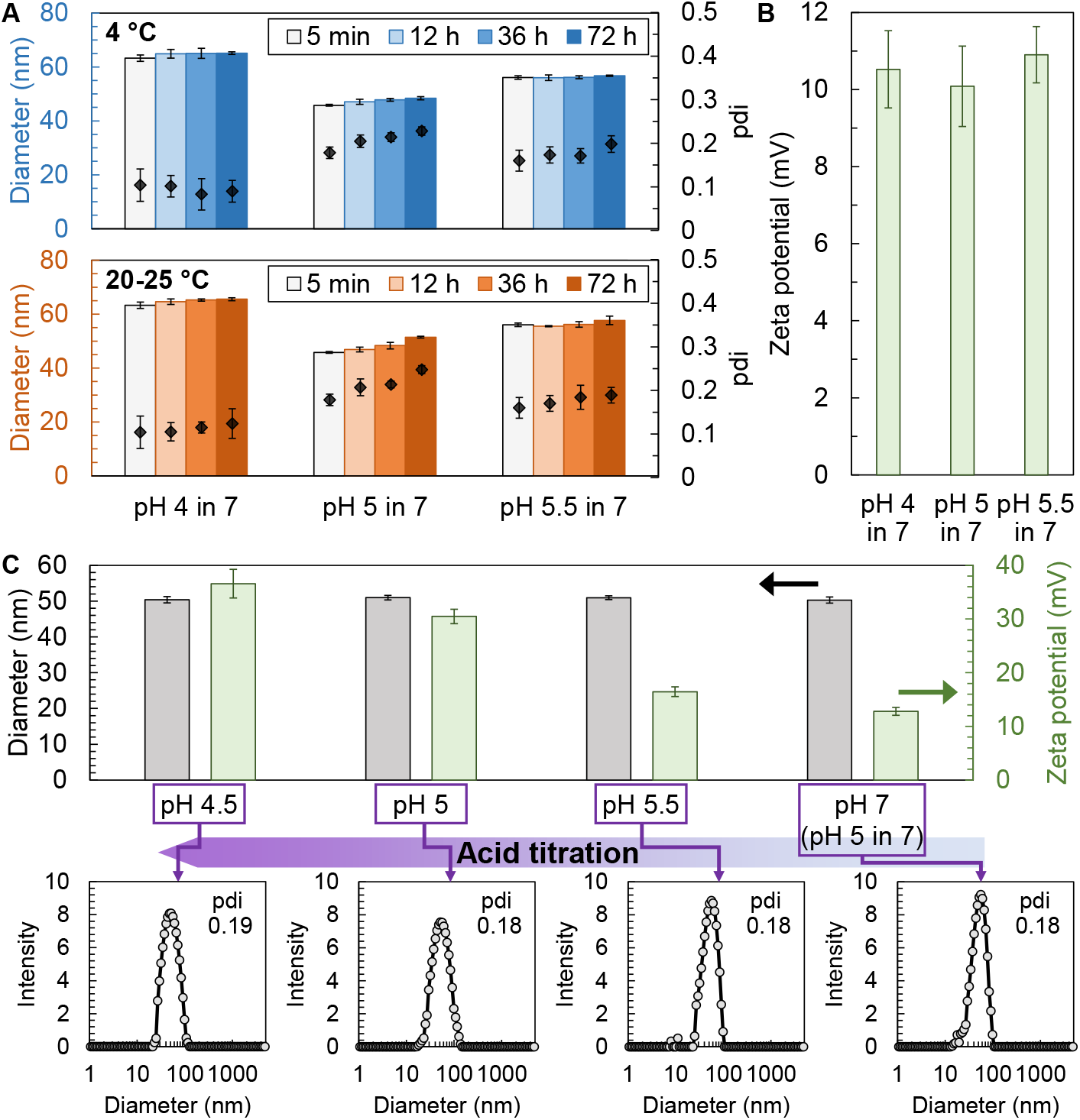
eLNP formulations. The hydrodynamic diameter (**A**) and zeta potential (**B**) of eLNPs measured by dynamic light scattering (DLS). eLNPs were made by mixing the ethanol solution of lipids with a buffer at pH 4, 5, or 5.5 in a two-inlet CIJ mixer and quenched the mixture into a pH 7 buffer, ensuring that the final dispersion contained 5 vol.% ethanol. The size of eLNPs was measured over time up to 72 h while keeping the samples refrigerated at 4 °C (top panel in A) and at room temperature (bottom panel in A). The diamonds in A show the averaged pdi for the corresponding sample. The zeta potential in B was measured within 5 min of preparation. **C**. The size and zeta potential of eLNPs that were prepared by mixing at pH 5 and quenching into a pH 7 buffer. The freshly made eLNPs were at a final pH of 7±0.1. The pH was adjusted to 5.5, 5, 4.5 by adding the appropriate amount of 100 mM acetate buffer at pH 4. The corresponding intensity-averaged size distribution and pdi of each case are shown.

#### Post-loading LNPs

We post-loaded yeast RNA (<170 nt), dasher green fluorescent protein mRNA (GFP mRNA, 1050 nt), Luciferase mRNA (Luc RNA, 1945 nt), or carrier RNA (2k-10k nt) (**Figure 3**) by adding the appropriate amount of RNA () in a pH 7 buffer to a quenched eLNP dispersion, followed by titration to pH 5.5. Mixing at neutral pH followed by protonation avoided aggregation, which was observed if post-loading was attempted directly at low pH. The quantity of added RNA was adjusted to achieve an N:P=6, which is optimal for in vivo transfections.^55–57^ Figure S2A (in the Supporting Information) shows that there is a rapid increase in the hydrodynamic diameter of LNPs after post-loading from approximately 55 nm for eLNPs (**Figure 2**) to approximately 95 nm for both yeast and carrier RNA post-loaded LNPs with colloidal stability over extended hours at room temperature. This indicates spontaneous loading independent of the RNA payload.

**Figure 3.**
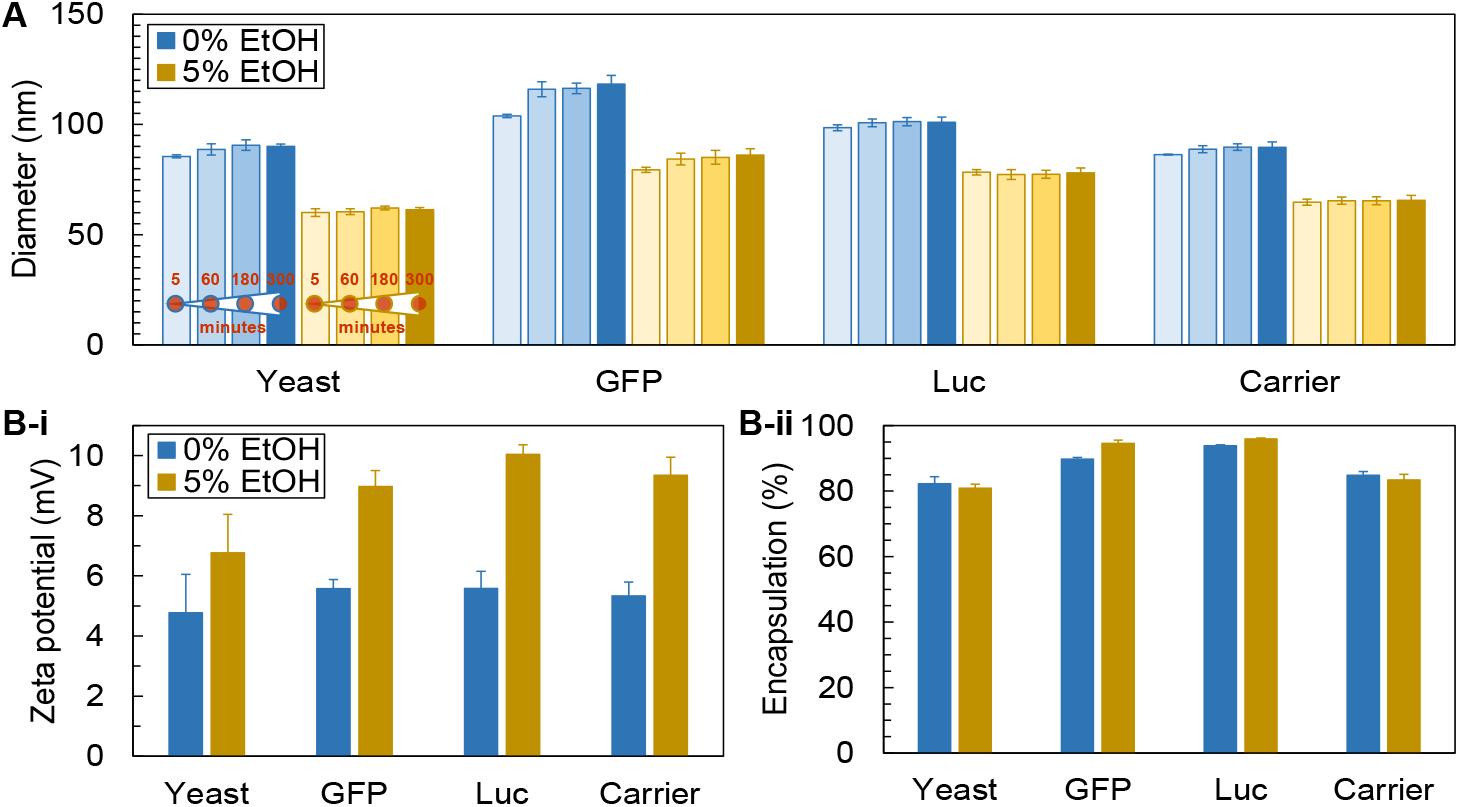
Post-loading LNPs with different RNAs. **A**. The averaged hydrodynamic diameter of LNPs post-loaded with four different RNAs: yeast RNA (<170 nt), GFP mRNA (1050 nt), Luc mRNA (1945 nt), and carrier RNA (2k-10k nt) at two ethanol content levels of 0 and 5 vol.%. The measurements were conducted for up to 5 hours for each sample. The averaged zeta potential (**B-i**) and encapsulation efficiency (**B-ii**) of post-loaded LNPs in A after 300 minutes of incubation at two different levels of ethanol. In all of these experiments, eLNPs were formulated at pH 5 and quenched at pH 7 (**Figure 2**) and post-loading was conducted at pH 5.5. Error bars are the standard deviation of three replicates.

Protonation of eLNPs is a crucial step in the post-loading process. When post-loading occurs at a neutral pH, there are insufficient electrostatic or other driving forces to effectively attract and adsorb free RNA onto eLNPs. As a result, most RNA only adsorbs to the eLNP surface, leading to a low EE of less than 50% (see Figure S3B in the Supporting Information). Surface-bound RNA at neutral pH also resulted in a much lower zeta potential compared to post-loading at pH 5.5 (see Figures S2A and S3A in the Supporting Information). Importantly, when larger RNAs (e.g., carrier RNA) were post-loaded at neutral pH, the surface-adsorbed RNA acted as a flocculant, causing colloidal instability in the LNPs. Consequently, the post-loaded LNPs exhibited over a 40% increase in size, from approximately 77 nm to 110 nm within 9 hours.

Three indicators were utilized to assess the internalization of RNA into the core of the LNPs, rather than adsorbed on the surface, when post-loaded at pH 5.5. First, positive zeta potentials of post-loaded LNPs (see Figure S2A in the Supporting Information) indicate internalization of the added RNA into the eLNP dispersion. Second, EEs of 80-90% were achieved for both yeast and carrier RNAs within 30 minutes of post-loading (see Figures S2B in the Supporting Information), which is equivalent to EEs observed for co-loaded LNPs. The EE is higher for the larger RNA. Third, the synchrotron scattering (discussed below) shows internal structures due to internalized RNA, that is equivalent to the scattering for co-loaded LNPs.^62^

Besides pH, we investigated the effect of ethanol concentration on post-loading by adding ethanol as well as by removing it from eLNP dispersions via rotary evaporation at room temperature under 30 Torr for approximately 1 hour, followed by filtration using 100 nm polyethersulfone (PES) syringe filters. It is expected that ethanol could fluidize the core of the LNP, affecting internalization of the RNA. Ethanol removal resulted in an increase in size, which may be due to lipid rearrangement.^63^ In our studies, the hydrodynamic diameter of eLNPs increased from 46 nm to 64 nm following ethanol removal. We subsequently post-loaded eLNPs with four different RNAs: yeast, Luc, GFP, and carrier RNAs, all at N:P=6. **Figure 3A** compares the hydrodynamic diameters of post-loaded LNPs containing these RNAs with and without ethanol over time. LNPs post-loaded at 0% ethanol retained similar stability to those post-loaded at 5% ethanol, with only a slightly greater size increase over 5 h. Post-loading at 10% and 15% ethanol concentrations yielded similar behavior to that at 5% (see Figure S4A in the Supporting Information). Lower zeta potentials of post-loaded LNPs at 0% ethanol suggest reduced lipid mobility in the core that may result in less RNA internalization (**Figure 3B-i**). However, the RNA assays indicated similarly high EE for all ethanol contents, reflecting low free RNA in the media (**Figure 3B-ii** and Figure S4B in the Supporting Information). To assess the internal structure of the LNPs, we conducted small-angle x-ray scattering (SAXS) and small-angle neutron scattering (SANS) of co-loaded and post-loaded LNPs.

### Post-loading mechanism of LNPs

The cryo-EM images in **Figure 4A** show that both eLNPs and post-loaded LNPs are spherical with electron-dense cores, supporting their suitability for core-shell models. **Figure 4B** compares SAXS patterns of eLNPs with both co-loaded and post-loaded yeast RNA LNPs. Scattering from eLNPs is consistent with polydisperse spherical particles with no detectable internal structure, as shown in cryo-EM images in **Figure 4A**. Nonetheless, the addition of RNA by post-loading or co-loading introduces a scattering peak near *q* ∼ 0.1 A°^-1^, indicating the development of similar internal structure with a correlation distance (ξ) of 6.3 nm 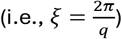.^49^

**Figure 4.**
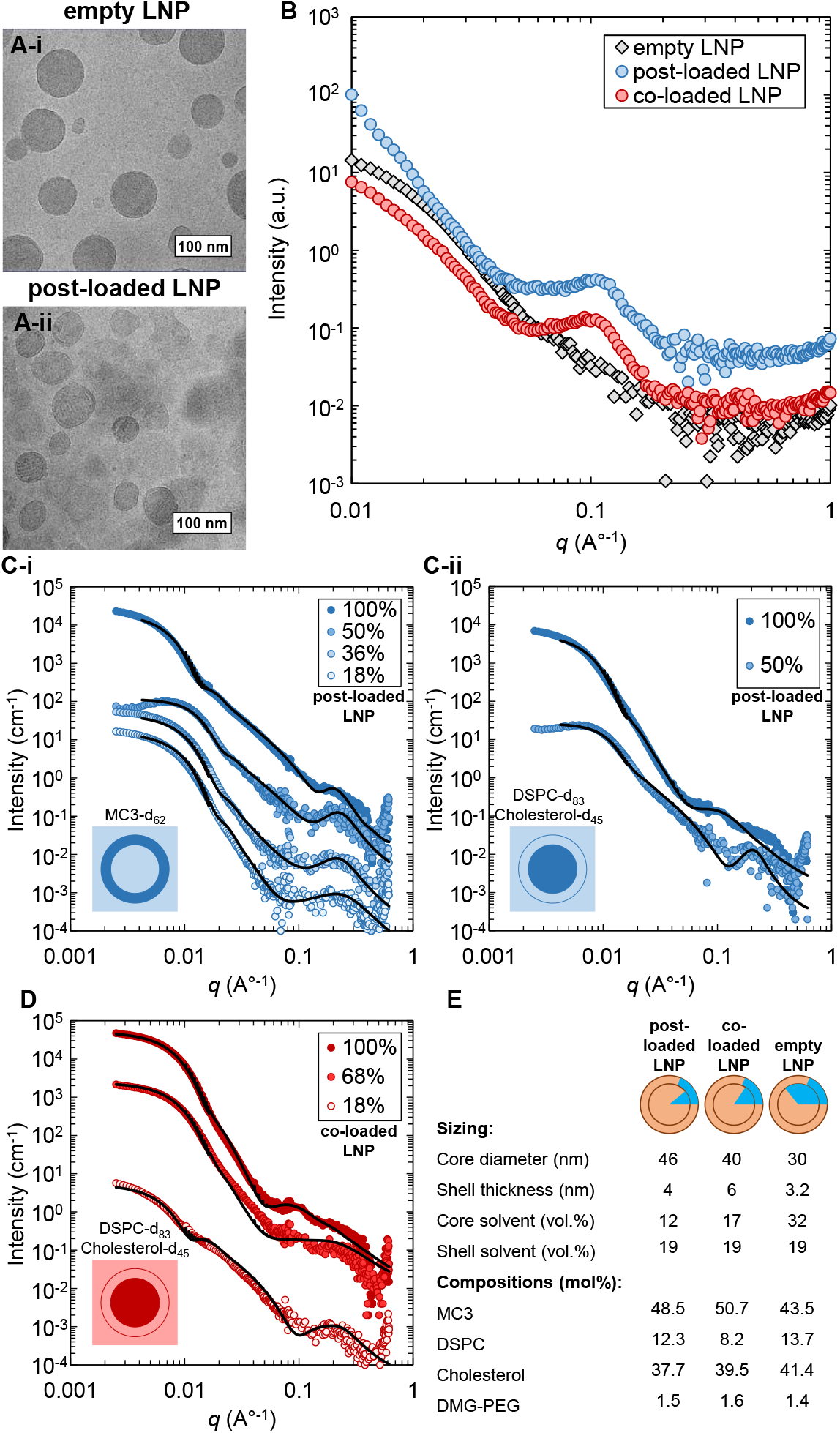
Structural analysis of LNPs. **A**. Cryo-EM images of empty LNPs (eLNPs, A-i) and post-loaded LNPs (A-ii). eLNPs were fabricated by mixing at pH5 and quenching at pH 7 (see Materials and Methods for details). Post-loaded LNPs were obtained after the addition of yeast RNA to eLNP (shown in A-i) followed by titrating the pH down to 5.5. **B**. SAXS data for the eLNPs and post-loaded LNPs shown in A, as well as co-loaded LNPs. SANS data for post-loaded (**C**) and co-loaded (**D**) LNPs at varying D_2_O volumetric fractions as noted in the legend. In **C-i** and **D**, formulations included deuterated MC3 (MC3-d_62_), whereas in **C-ii**, cholesterol and DSPC were replaced with deuterated cholesterol (cholesterol-d_45_) and DSPC (DSPC-d_83_), respectively. This setup enables the analysis of the shell and core structures independently. Solid lines represent core-shell model fits to the data, as described in the manuscript. **E**. The structural and compositional analyses of post-loaded, co-loaded, and empty LNPs. The corresponding cartoons represent the volume fraction of solvent (in blue) at the core and in the shell of LNPs.

The internal structure of LNPs was determined by internal contrast variation in SANS experiments, achieved by selective deuteration of lipid components (see Table S1 in the Supporting Information); one formulation contained deuterated MC3 (MC3-d_62_) while the other contained deuterated cholesterol (cholesterol-d_45_) and DSPC (DSPC-d_83_) with other lipid components hydrogenated (and RNA for loaded LNPs). These LNPs were dispersed in the mixtures of D_2_O and H_2_O buffers at different volume fractions. This yielded 6-10 different H/D isotopic compositions, enabling a higher resolution of LNP internal structure. **Figure 4C-i** and **C-ii** show SANS measurements for post-loaded LNPs with yeast RNA at pH 5 where eLNPs were formulated by mixing at pH 4 and quenching in pH 7.

At 100% D_2_O, no specific features were observed, whereas at 50% D_2_O a broad peak at *q* ∼ 0.19 A°^-1^ appeared, with the expected contrast match point at 50-68% D_2_O. At the contrast match point of the entire LNP, the overall scattering is minimized, causing the appearance of the broad feature because the nanoscale scattering is maximized. Importantly, this peak appeared for co-loaded LNPs as well (see **Figure 4D**) in line with the interpretation of SAXS measurements. We found a core-shell model that incorporates broad peak best fit our data.^64^

To examine the internal consistency, we computed the mole fraction of each lipid component from the contrast for each component and then compared them to the expected values used in the formulation (**Figure 4E**). In this approach, the water content of the shell is independent of the mRNA internalization by co- or post-loading and remains constant at 19 vol.% water (see **Figure 4E**). Additionally, **Figure 4E** shows that the core contained 17 vol.% water for co-loaded LNPs formulated when mixed at pH 4 and quenched into a pH 7 bath. The fitting of post-loaded LNPs indicated 12 vol.% water in the core which is in reasonable agreement to co-loaded LNPs.

For eLNPs with no RNA at the core, the core contained 65 vol.% water (**Figure 4E** and Figure S5 in the Supporting Information) under the same pH formation and quenching conditions. For eLNPs, there is excess cationic charge in the core since none of the ionized lipids are condensed with RNA ribose sugars. The condensation both minimizes the waters of hydration associated with ionized lipids and creates compressive elastic stresses that condense some of the aqueous volumes in the core.

### Transfection and cytotoxicity of post-loaded LNPs

To demonstrate that the structural and physical chemistry similarities between co-loaded and post-loaded LNPs translate to comparable performance, we evaluated their transfection efficacy and cytotoxicity. Our assessments were conducted using HeLa cells as the model system. We post-loaded LNPs with GFP mRNA. After varying the loading time between 30 min to 6 hours, we introduced the post-loaded LNPs to the cell cultures for a total of 24 hours of incubation at 37 °C. While the physical chemistry experiments indicated post-loading was rapid (Figure S2 in the Supporting Information), we need to determine whether there was a required loading time to reach maximum transfection efficiency. **Figure 5A** illustrates that after 30 minutes of post-loading, the highest transfection to the cells is observed, persisting for at least 4 hours before gradually declining, likely due to RNA degradation. In a clinical application, the administration within four hours would be feasible.

**Figure 5.**
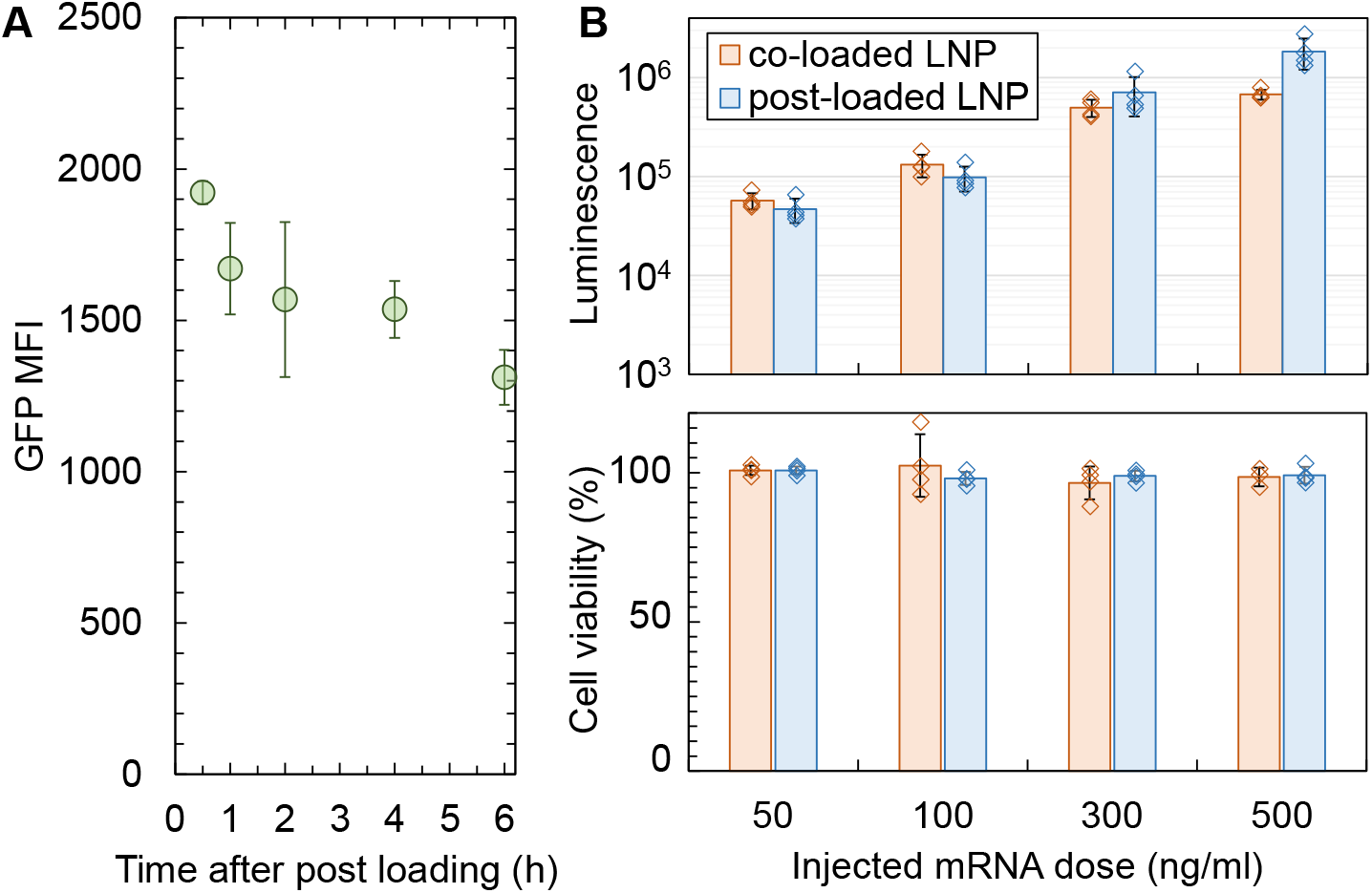
mRNA transfections to HeLa cells. **A**. The mean fluorescence intensity (MFI) of GFP mRNA delivered to HeLa cells by post-loaded LNPs (PNLPs). PLNPs were prepared by formulating eLNPs at pH 5 and quenching at pH 7 (**Figure 2**), followed by post-loading with GFP mRNA at pH 5.5 for various loading times, as indicated on the x-axis. HeLa cells were treated with PLNPs containing 500 ng/ml GFP mRNA in the well and then incubated at 37 °C and 5% CO_2_ for 24 hours. Subsequently, green protein expression was measured four times using a plate reader, with error bars representing the standard deviation of these four readouts. **B**. The averaged luciferase activity and viability percentage of HeLa cells treated with co-loaded and post-loaded LNPs dosed at 50, 100, 300, and 500 ng/ml Luc mRNA in the well (bars). For each sample, luciferase fluorescence intensity was measured four times (symbols), and error bars report the standard deviation of these four readouts. All these LNP formulations were at 5 vol.% ethanol.

We formulated post-loaded LNPs and co-loaded LNPs with Luc mRNA at pH 5.5. These formulations exhibited similar physicochemical properties, including size, zeta potential, and EE (see Figure S6A in the Supporting Information). **Figure 5B** reveals that the dose-response data indicate that the co-loaded and post-loaded LNPs behave equivalently. We repeated the post-loading using eLNPs prepared at pH 4 and quenched at pH 7 as well as for eLNPs prepared at pH 5 and quenched at pH 7, labelled as PLNP1 and PLNP2 on Figure S6 (in the Supporting Information), respectively. Figure S6B shows that these two eLNP formulations achieved comparable transfection efficiencies, with no appreciable difference from the co-loaded LNPs.

## Conclusions

The electrostatic binding between messenger ribonucleic acid (mRNA) and cationic ionizable lipid is the key step during their co-precipitation to form lipid nanoparticles (LNPs) that contain encapsulated mRNA (co-loaded LNPs). Instead of relying on this traditional co-loading method, we demonstrated how the electrostatic complexation between mRNA and ionizable lipid can be tuned in “empty” LNPs (eLNPs) to establish a “post-loading” protocol. In this approach, colloidally stable eLNPs are first produced during the rapid mixing of lipid solution with aqueous buffer using confined impinging jet (CIJ) technology. Subsequently, RNA is introduced to the eLNP suspension, and the pH is titrated from neutral pH down to one unit below the pKa of the ionizable lipid while maintaining ethanol concentration to ensure lipid mobility without compromising LNP stability. Both co-loading and post-loading routes yield equivalent LNPs, as determined by four separate experimental assays. First, LNPs with a hydrodynamic diameter of <100 nm and encapsulation efficiency of 80-95% were synthesized through both co-loading and post-loading routes. The zeta potentials of both LNPs are similar (approximately +8 mV), indicating that significant negatively charged RNA does not reside at the surface of LNPs. Second, synchrotron x-ray scattering indicates the same internal structure (with a scattering peak at *q* ∼ 0.1 A°^-1^ and with similar peak intensities). Third, neutron scattering shows similar core and shell compositions. Finally, mRNA-loaded LNPs formulated through the post-loading route exhibit similar protein expression levels to those produced via the co-loading route. These promising results offer a more flexible and adaptable approach for designing vaccines, meeting the dynamic needs of global health crises. In addition, the results indicate that eLNPs can be produced to be used in laboratory research settings to evaluate mRNA sequences and delivery. eLNPs can be used as a replacement of the cationic Lipofectamine® reagents that are often used, but which cannot be translated to clinical formulations, whereas the eLNP components are those used in FDA-approved vaccine formulations.

## Author contributions

N.B. and R.K.P. conceptualized the experiments; N.B., D.F.A., and S.N. performed and analyzed formulation experiments; N.B. and S.N. designed, performed and analyzed the small-angle x-ray scattering experiments; N.B., S.N.S., and B.K. performed and analyzed cell transfection experiments; D.F.A., L.S.M., K.W., and G.G.W. designed, performed, and analyzed the small-angle neutron scattering experiments; N.R.Y., M.M., and M.C. synthesized and characterized the deuterated lipids; N.Z. processed and analyzed cryo-EM images; N.B., G.G.W., and R.K.P. wrote, reviewed, and edited the original draft, and R.K.P. funded and supervised the overall project.

## Data availability

A data availability statement (DAS) is required to be submitted alongside all articles. Please read our [full guidance on data availability statements](https://www.rsc.org/journals-books-databases/author-and-reviewer-hub/authors-information/prepare-and-format/data-sharing/#dataavailabilitystatements) for more details and examples of suitable statements you can use.

## Acknowledgements

We greatly acknowledge the support of the lead scientist Lin Yang and beamline scientist Shirish Chodankar at Brookhaven National Laboratory. The LiX beamline is part of the Center for BioMolecular Structure (CBMS), which is primarily supported by the National Institutes of Health, National Institute of General Medical Sciences (NIGMS) through a P30 Grant (P30GM133893), and by the DOE Office of Biological and Environmental Research (KP1605010). LiX also received additional support from NIH Grant S10 OD012331. As part of NSLS-II, a national user facility at Brookhaven National Laboratory, work performed at the CBMS is supported in part by the U.S. Department of Energy, Office of Science, Office of Basic Energy Sciences Program under contract number DE-SC0012704. We also acknowledge the support of the Australian Government in provision of access to ANSTO’s National Deuteration Facility and Australian Centre for Neutron Scattering, which are partly funded through the National Collaborative Research Infrastructure Strategy (NCRIS), via proposal D/N15402, and the support of SANS beamline scientist Kathleen Wood. We appreciate support from the FDA under contract award 75F40122C001 to Princeton University. We acknowledge the support from Tessera Therapeutics Inc. for initial funding on the project. Also, we appreciate support from Karthik Nagapudi of Genentech to enable the TEM analysis through the University of Wisconsin. Cryo-EM imaging was performed in the cryo-EM Research Center (CEMRC) in the Department of Biochemistry at the University of Wisconsin-Madison. We also acknowledge Bill and Melinda Gates Foundation under INV-056149 for RKP.

