## Supplementary material for "Post-loading ribonucleic acid into lipid nanoparticle carriers": SI

**This PDF file includes:**

Figures S1 to S14
Schemes S1 to S2
Table S1
Computations for Debye length, PEG brush thickness, synthesis and characterization of deuterated lipids, and SANS data analysis

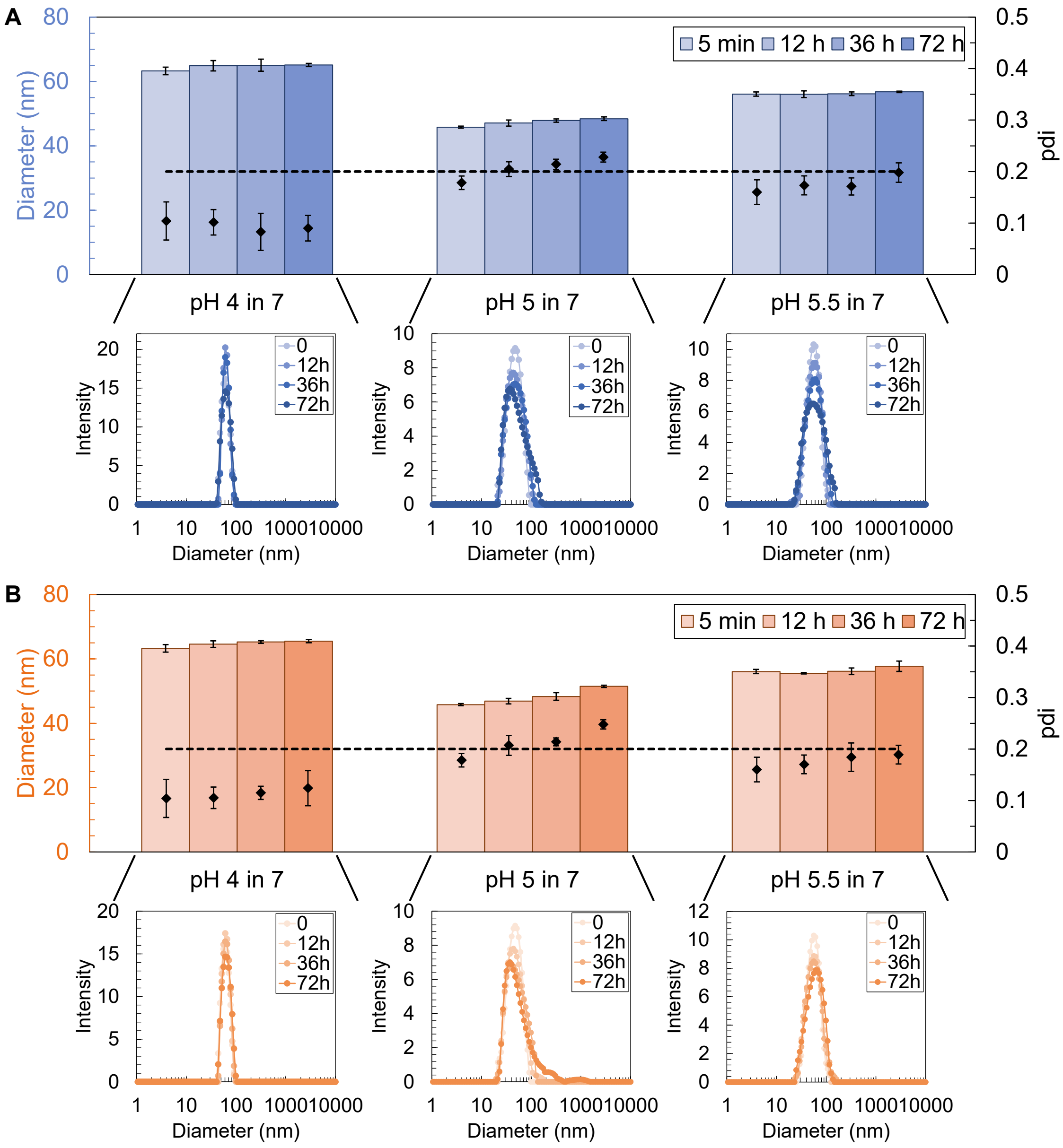


**Figure S1. Stability of empty lipid nanoparticles (eLNPs).** The changes in the mean hydrodynamic diameter and the corresponding size distribution of eLNPs over time stored at 4 °C (**A**) and at room temperature (20-25 °C) (**B**). The eLNPs were made using CIJ mixers under the marked mixing and quenching pH (see Figures 1B in the main manuscript for the reference). The diamond symbols in A and B represent averaged pdi for the corresponding case and the dashed line shows the critical pdi 0.2.

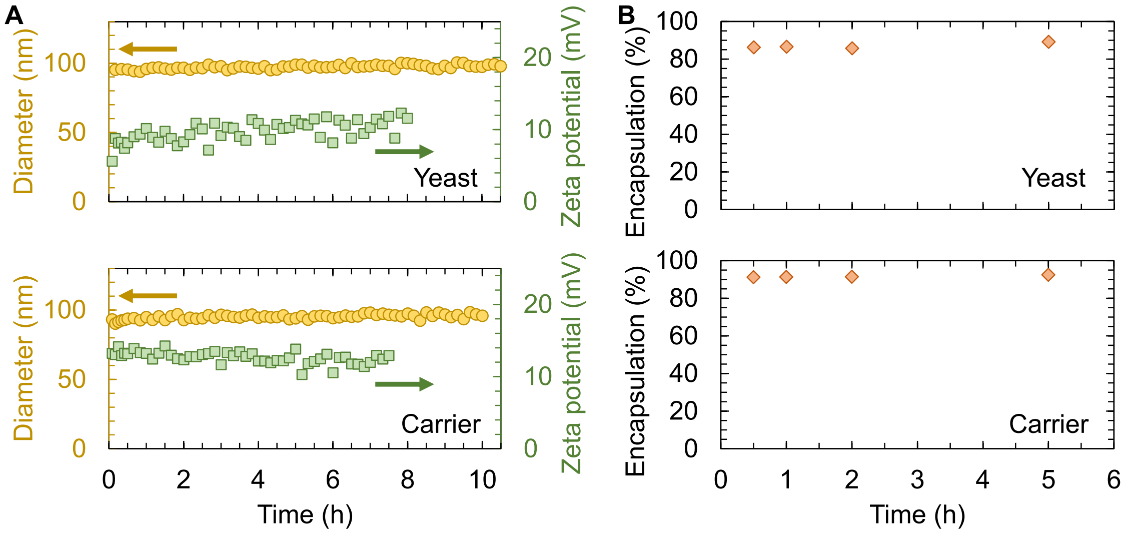


**Figure S2. Post-loading kinetics at pH 5. A.** The size and zeta potential measurements of LNPs post-loaded with yeast (top panel) and carrier (bottom panel) RNA. These measurements were conducted over time during which eLNPs and added RNA were incubated at room temperature (i.e., post-loading time). eLNPs were made by mixing lipid solution with acetate buffer at pH 5 and quenching into a pH 7 HEPES buffer and the post-loading step was conducted at pH 5.5 (see the Materials and Methods in the manuscript for details). **B.** The encapsulation efficiency of yeast (top panel) and carrier (bottom panel) RNA post-loaded into LNPs as a function of post-loading time.

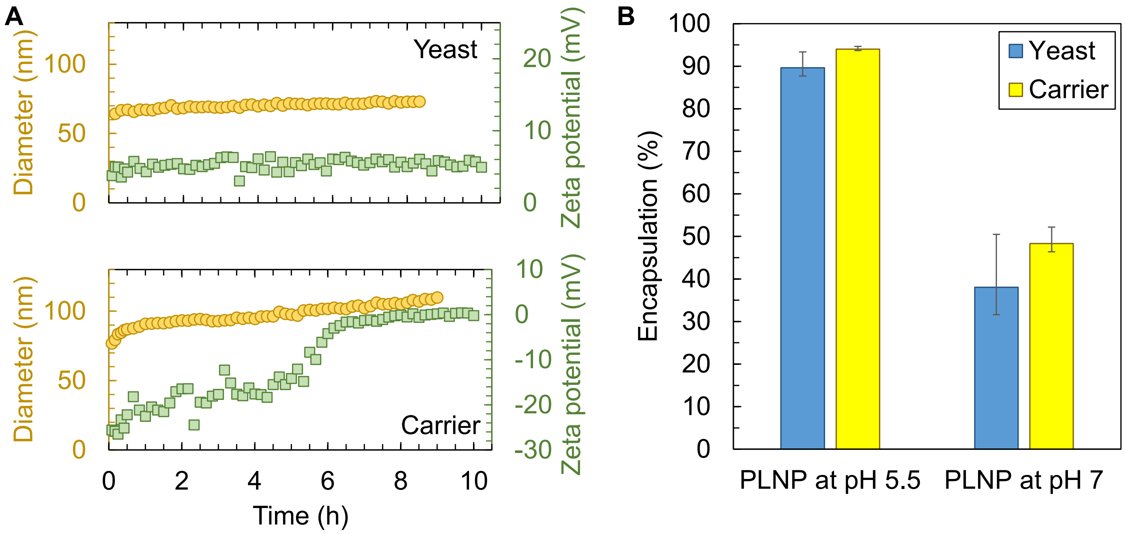


**Figure S3. Post-loading kinetics at pH 7. A.** The size and zeta potential measurements of LNPs post-loaded with yeast (top panel) and carrier (bottom panel) RNA. These measurements were conducted over time during which eLNPs and added RNA were incubated at room temperature (i.e., post-loading time). eLNPs were made by mixing lipid solution with acetate buffer at pH 5 and quenching into a pH 7 HEPES buffer and the post-loading step was conducted at pH 7. **B.** The encapsulation efficiency for post-loaded LNPs at pH 7 in comparison to that at pH 5.5.

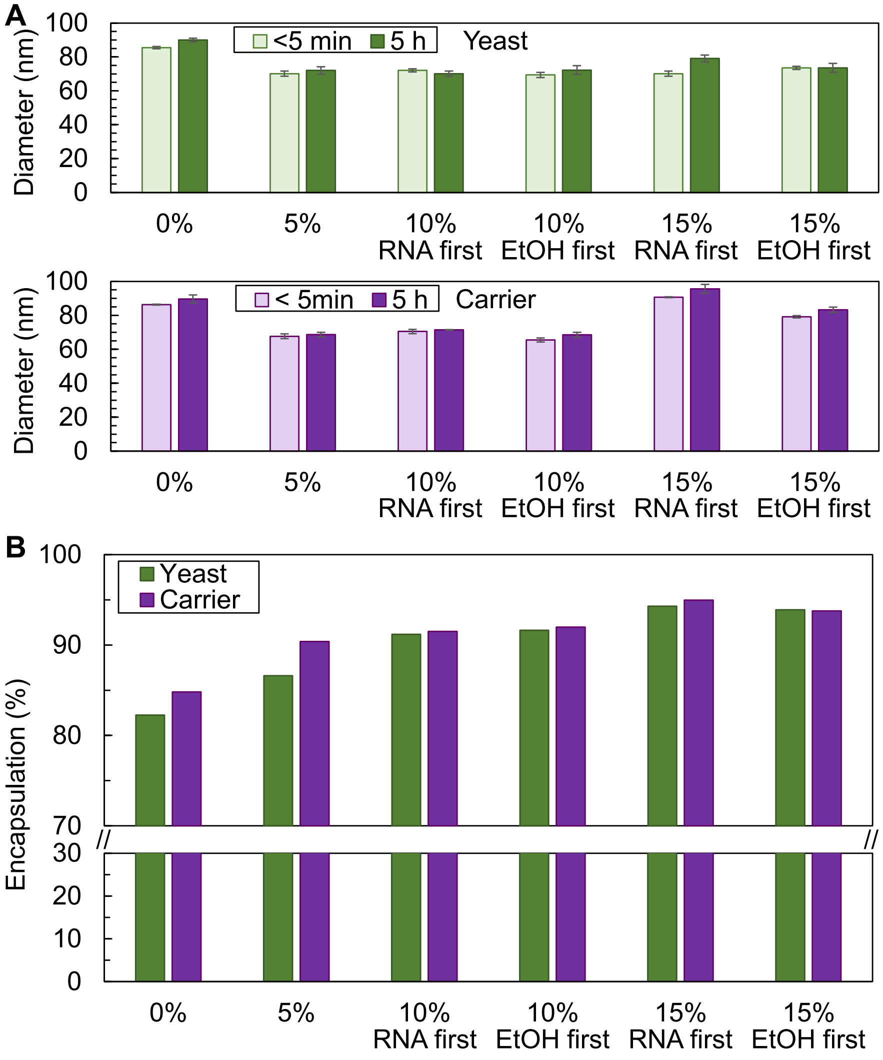


**Figure S4. The ethanol effect on post loading capacity.** Size (**A**) and encapsulation efficiency (**B**) of yeast and carrier RNA post-loaded in eLNPs as a function of ethanol content. eLNPs were made by mixing lipid solution with acetate buffer at pH 5 and quenching them into a pH 7 buffer and the post-loading step was conducted at pH 5.5 (see the Materials and Methods in the manuscript for details). The final ethanol content in these samples was 5 vol.%. For post-loading at 0%, ethanol was removed *via* rotary evaporation at room temperature (see the manuscript for details). For post-loading at higher ethanol contents of 10 vol.% and 15 vol.%, two routes were examined: adding RNA first followed by ethanol to achieve 10 vol.% or 15 vol.% (labeled as "RNA first") or adjusting the ethanol content to 10 vol.% or 15 vol.% first, followed by the addition of RNA (labeled as "EtOH first").

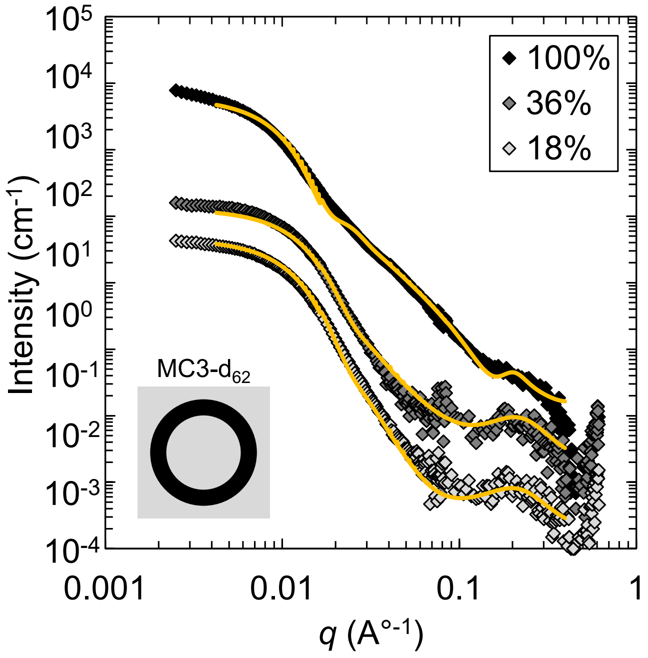


**Figure S5. SANS data for eLNPs.** SANS data for empty LNPs (eLNPs) at varying $D_2O$ volumetric fractions as noted in the legend. eLNPs formulations included deuterated MC3 (MC3-$d_{62}$).

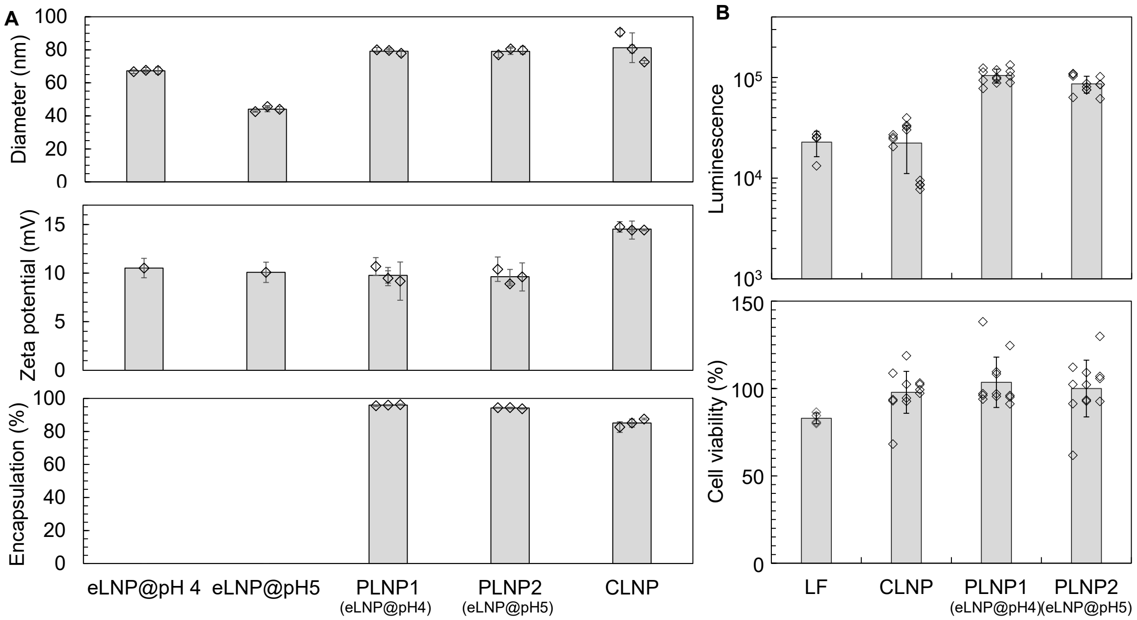


**Figure S6. Physicochemical characterizations and transfections of different formulated LNPs. A.** The averaged size (top panel), zeta potential (middle panel), and encapsulation efficiency (bottom panel) of LNPs formulated for cell transfection studies shown in Figure 5 of the manuscript. Co-loaded LNPs (CLNPs) were prepared with the current co-precipitation technology at pH 5.5, while post-loaded LNPs (PLNPs) were prepared by post-loading eLNPs at pH 5.5. Empty LNPs (eLNPs) were prepared by mixing either at pH 4 or pH 5. Three samples were examined for loaded LNPs (symbols), each of which is the average of three readings. **B.** The averaged luciferase activity (top panel) and viability percentage (bottom panel) of HeLa cells treated with Lipofectamine 3000 (LF), CLNPs, and PLNPs (bars). Luciferase fluorescence intensity was measured four times for each sample (symbols) and three samples were examined for both CLNPs and PLNPs. Error bars represent the standard deviation of the pooled data. Luc mRNA was used in all loaded LNPs, with HeLa cells dosed at 500 ng/ml mRNA in the well.

**Debye length and PEG brush thickness calculations**

The Debye length ( $\kappa^{-1}$ ) is calculated from $\kappa^{-1} = \sqrt{\frac{\varepsilon_0 \varepsilon_r k_B T}{\sum_i (z_i e)^2 C_i}}$ (1), where $\varepsilon_0$ and $\varepsilon_r$ are vacuum permittivity (i.e., $8.85 \times 10^{-12}$ F/m) and relative dielectric constant of the solvent, respectively, $k_B$ is Boltzmann constant (i.e., $1.38 \times 10^{-23}$ $m^2kg/s^2K$), $T$ is absolute temperature, $e$ is the elementary charge (i.e., $1.602 \times 10^{-19}$ C), $z$ is the valance charge of ion $i$ , and $C_i$ is the number concentration of ion $i$ (in $1/m^3$). Assuming that HEPES buffer is the dominant at ~10 mM and neglecting the 5 vol.% of ethanol on $\varepsilon_r$ , we compute $\kappa^{-1} \cong 3$ nm. Given the low grafting density of PEG on the LNP surface, we expect a mushroom regime for the PEG layer with a 3.6 nm thickness (i.e., $2R_g$ ) estimated from $2R_g = 0.043 M_W^{0.583}$ , where $M_W$ is the molecular weight of PEG in g/mol and $R_g$ is radius of gyration in nm (2).

**Synthesis of Deuterated Materials**

*General Experimental Synthesis*

All reactions were performed under an atmosphere of nitrogen unless otherwise specified. Chemicals and reagents of the highest grade were purchased from Sigma-Aldrich/Merck and were used without further purification. Solvents were purchased from Sigma-Aldrich and Merck. NMR solvents were purchased from Cambridge Isotope Laboratories Inc. (MA, USA) and Sigma-Aldrich (Merck) and were used without further purification. $D_2O$ (99.8%) was supplied Merck. Anhydrous dichloromethane, tetrahydrofuran and diethyl ether were obtained from an LC Technology Solutions Inc. SP-1 Stand Alone Solvent Purification System. Analytical thin-layer chromatography (TLC) was performed using Merck aluminium backed silica gel 60 F254 (0.2 mm) plates, which were visualised with shortwave (254 nm) ultraviolet light or with potassium permanganate, vanillin, Hanessian's or bromocresol green stains. Flash column chromatography was performed using a Buchi Pure automated system, with the eluent mixture reported as the volume:volume ratio. Hydrothermal reactions were performed in $D_2O$ at high temperatures in a Mini Benchtop 4560 Parr reactor (600 ml vessel capacity, 3000 psi maximum pressure, 350 °C maximum temperature) (Moline, USA). Nuclear magnetic resonance spectra were recorded at 300 K using a Bruker AVANCE DRX400 (400 MHz) spectrometer. $^1H$ chemical shifts are expressed as parts per million (ppm) with residual chloroform (δ 7.26) as reference and are reported as chemical shift (δ); relative integral; multiplicity; coupling constants (*J*) reported in Hz. Assignments in $^1H$ NMR are quoted as residual H except where the carbon is fully protiated. $^{13}C$ chemical shifts are expressed as parts per million (ppm) with residual chloroform (δ 77.16) as reference and reported as chemical shift (δ); multiplicity. $^{13}C$ resonances attached to deuterium appear as multiplets when only the proton nucleus is decoupled $^{13}C$ {$^1H$} and resolve to singlets when both proton and deuterium nuclei are decoupled (i.e., $^{13}C$ {$^1H$, $^2H$}). $^2H$ chemical shifts are reported as parts per million (ppm) and are reported as chemical shift (δ); multiplicity. $^{31}P$ chemical shifts are reported as parts per million (ppm) and are reported as chemical shift (δ). High-Resolution mass spectrometry was performed using a Shimadzu 9050 Time-of-flight spectrometer hyphenated to a Shimadzu 40 series UHPLC system. Samples (0.5 µL) were injected directly to the ESI source in a mobile phase of 80:20 ACN:$H_2O$ with 0.1% acetic acid as an ionisation aid at a flow rate of 0.4 ml/min. M/Z profile data was exported and analysed using DGet! to obtain the overall deuteration levels and the distribution of isotopologues (3).

*Details of Synthesis*
1,2-Distearoyl-$d_{70}$-sn-glycero-3-phosphocholine-$d_{13}$ (**DSPC-$d_{83}$ average 96%D**)

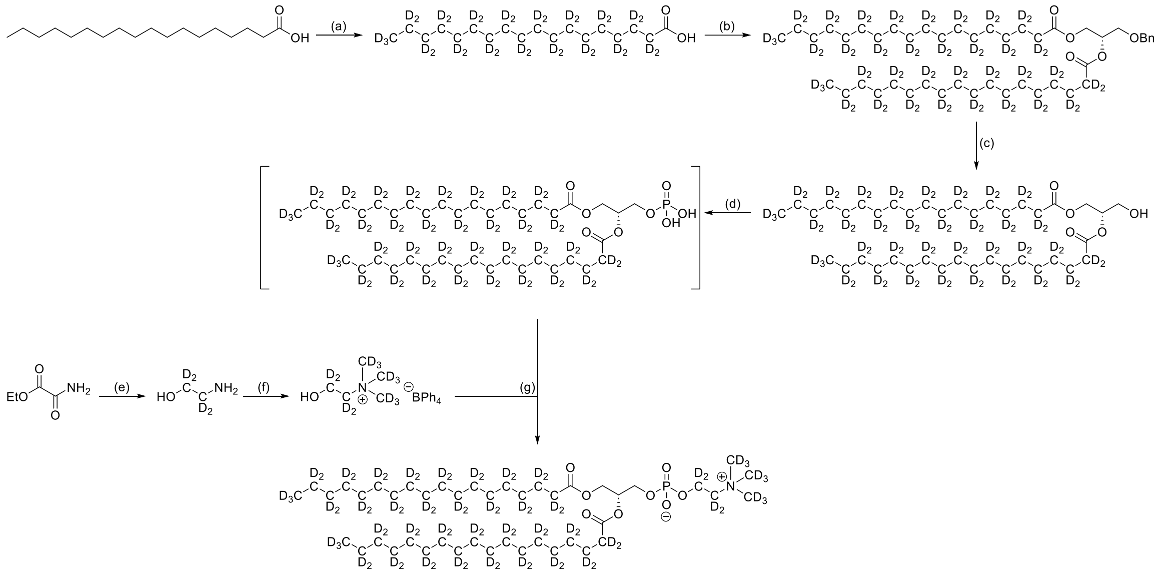

**Scheme S1. Synthesis steps of DSPC-$d_{83}$.** Reagents and conditions: (a) Pt/C, NaOH, $D_2O$, 220 °C, 3 days, 2 cycles; (b) 3-O-Benzyl-sn-glycerol, DCC, DMAP, $CH_2Cl_2$, room temperature, 24 h; (c) $H_2$, Pd/C, EtOH, THF, room temperature, 22 h; (d) i) $POCl_3$, $Et_3N$, $CH_2Cl_2$, 0 °C, 50 mins, ii) $H_2O$, room temperature, 1 h; (e) $LiAlD_4$, THF, reflux, 16 h; (f) i) NaOH, $CD_3I$, EtOH, $H_2O$, 0 °C to room temperature, 16 h, ii) $NaB(Ph)_4$, EtOH, $H_2O$, room temperature, 1.5 h; (g) Tripsyl chloride, pyridine, room temperature, 12 h.

<u>Stearic acid-$d_{35}$:</u>

Stearic acid (13 g, 46.7 mmol), sodium hydroxide (2.2 g, 56 mmol) and platinum on activated carbon (10% w/w, 0.6 g) were added to a 600 ml Parr reactor vessel and suspended in deuterium oxide (120 ml). The vessel was flushed with nitrogen and heated at 220 °C for 3 days. The mixture was cooled to room temperature, acidified with hydrochloric acid (5 M), diluted with ethyl acetate (100 ml) and stirred for 10 minutes. The biphasic mixture was filtered through a pad of Celite®. The filtrate was transferred to a separatory funnel and the layers separated. The aqueous phase was extracted with ethyl acetate (3 × 50 ml), the organic phases combined, dried over anhydrous magnesium sulfate and concentrated under reduced pressure, to obtain stearic acid-$d_{35}$. The material was subjected to another cycle using the same conditions as above to obtain the deuterated fatty acid (95.5%D by mass spectrometry, quant.) as a white solid, $^1$H NMR (400 MHz, $CDCl_3$) δ 2.31 (bs, residual), 1.58 (bs, residual), 1.32-1.12 (m, residual), 0.82 (bs, residual) ppm; $^2$H NMR (61.4 MHz, $CDCl_3$) δ 2.32 (2D, bs), 1.59 (2D, bs), 1.42-0.98 (28D, bm), 0.84 (3D, s) ppm; $^{13}$C{$^1$H, $^2$H} NMR (100.6 MHz, $CDCl_3$) δ 179.7, 33.3, 30.8, 29.2-27.6 (m), 23.8, 21.6, 13.2 ppm.

<u>(*S*)-3-(Benzyloxy)propane-1,2-diyl bis(octadecanoate-$d_{35}$):</u>

To a solution of 3-O-benzyl-sn-glycerol (5.0 g, 27.5 mmol), stearic acid-$d_{35}$ (19.3 g, 60.4 mmol) and 4-dimethylaminopyridine (3.4 g, 27 mmol) in dichloromethane (300 ml) was added a solution of N,N'-dicyclohexylcarbodiimide (12.5 g, 60.5 mmol) in dichloromethane (20 ml). The mixture was stirred for 24 hours at room temperature and filtered to remove solids. The filtrate was then evaporated under reduced pressure to give a pale yellow residue, which was purified by flash chromatography using ethyl acetate, hexane (1:19) as an eluent to give the title compound (15 g,

70%) as a white solid, $^{1}H$ NMR (400 MHz, $CDCl_3$) δ 7.45-7.20 (5H, m), 5.38-5.18 (1H, m), 4.66-4.48 (2H, m), 4.44-4.31 (1H, m), 4.28-4.13 (1H, m), 3.70-3.54 (2H, m), 2.27 (bs, residual), 1.58 (bs, residual), 1.36-1.13 (bm, residual), 0.86 (bs, residual) ppm; $^{2}H$ NMR (61.4 MHz, $CDCl_3$) δ 2.27 (4D, bs), 1.56 (4D, bs), 1.45-0.97 (56D, bm), 0.84 (6D, s) ppm; $^{13}C\{^{1}H\}$ NMR (100.6 MHz, $CDCl_3$) δ 173.6, 173.3, 137.8, 128.5, 127.9, 127.7, 73.4, 70.1, 68.4, 62.8, 33.6 (m), 28.5 (m), 24.0 (m), 21.6 (m), 13.0 (m) ppm.

(*S*)-3-Hydroxypropane-1,2-diyl bis(octadecanoate-$d_{35}$):

To a solution of the benzyl ether (10 g, 12.7 mmol) in a mixed solvent of THF (90 ml) and EtOH (500 ml) at room temperature was added palladium on carbon (1 g, 10% w/w). The system was evacuated and backfilled with hydrogen three times and stirred at room temperature for 22 hours. The reaction mixture was filtered through Celite, and the filtrate concentrated. The crude residue was purified by flash column chromatography using ethyl acetate, hexane (1:9) as an eluent to afford the alcohol (3.2 g, 36%) as a white solid, $^{1}H$ NMR (400 MHz, $CDCl_3$) δ 5.08 (1H, quin. $J$ = 5.1 Hz), 4.32 (1H, dd, $J$ = 4.7, 12.3 Hz), 4.23 (1H, dd, $J$ = 5.7, 12.0 Hz), 3.77-3.68 (2H, m), 2.31 (bs, residual), 1.57 (bs, residual), 1.31-1.13 (bm, residual), 0.82 (bs, residual) ppm; $^{2}H$ NMR (61.4 MHz, $CDCl_3$) δ 2.31 (4D, bs), 1.58 (4D, bs), 1.44-0.95 (56D, bm), 0.83 (6D, s) ppm; $^{13}C\{^{1}H\}$ NMR (100.6 MHz, $CDCl_3$) δ 174.0, 173.7, 72.2, 62.1, 61.7, 33.5 (m), 28.5 (m), 24.2 (m), 21.5 (m), 13.1 (m) ppm.

(*R*)-3-(Phosphonooxy)propane-1,2-diyl bis(octadecanoate-$d_{35}$):

Triethylamine (0.67 g, 6.5 mmol) and 1,2-distearoyl-$d_{70}$-sn-glycerol (3 g, 4.3 mmol) in anhydrous dichloromethane (50 ml) were added to a solution of phosphorous oxychloride (0.8 g, 5.1 mmol) in dichloromethane (150 ml). The mixture was stirred for 50 minutes at 0 ˚C. Volatiles were removed to give a crude residue. Water (5 ml) was added to the crude, and the mixture stirred at room temperature for 60 minutes, and then evaporated to dryness. The crude phosphatidic acid (4 g) was used immediately without any further purification or characterisation.

Choline-$d_{13}$ tetraphenylborate:

Choline-$d_{13}$ tetraphenylborate was synthesised using previously reported methods (4).

1,2-Distearoyl-$d_{70}$-sn-glycero-3-phosphocholine-$d_{13}$

To the crude phosphatidic acid (4 g, 5.2 mmol) was added 2,4,6-triisopropylbenzenesulfonyl chloride (1.56 g, 5.2 mmol) and choline-$d_{13}$ tetraphenylborate (2.3 g, 5.2 mmol). The solids were dissolved in anhydrous pyridine (30 ml) and the reaction mixture was stirred for 12 hours at room temperature. The mixture was evaporated to dryness and the crude residue purified by flash column chromatography using water, methanol, dichloromethane 70:30:0.4 to give afford the phosphocholine (0.9 g, 33%) as a white solid, $^{1}H$ NMR (400 MHz, $CDCl_3$) δ 5.28-5.14 (1H, m), 4.38 (1H, dd, J = 3.1, 12.1 Hz), 4.23 (bm, residual), 4.12 (1H, dd, J = 7.3, 12.1 Hz), 4.05-3.91 (2H, m), 3.84 (bs, residual), 3.34 (bs, residual),2.26 (bm, residual), 1.53 (bs, residual), 1.32-1.11 (bm, residual), 0.82 (bs, residual) ppm; $^{2}H$ NMR (61.4 MHz, $CDCl_3$) δ 3.36 (9D, bs), 2.27 (4D, bs), 1.52 (4D, bs), 1.48-0.91 (56D, bm), 0.83 (6D, s) ppm note methylene positions of choline head-group were not resolved; $^{31}P$ NMR (161.9 MHz, $CDCl_3$) δ -1.13 ppm; $^{13}C\{^{1}H, ^{2}H\}$ NMR (100.6 MHz, $CDCl_3$) δ 173.7, 173.4, 70.5 (m), 65.2 (m), 63.6 (m), 63.0, 58.8 (m), 53.4, 33.6, 33.5, 30.7, 28.7-28.3 (m), 28.2, 28.2, 28.0, 28.0, 24.0, 23.9, 21.5, 13.1 ppm; MS (ESI) Calc. $C_{44}H_5D_{83}NO_8P$ $[M+H]^+$: 874.1536. Deuteration level calculated as 96.5% using DGet! (3).

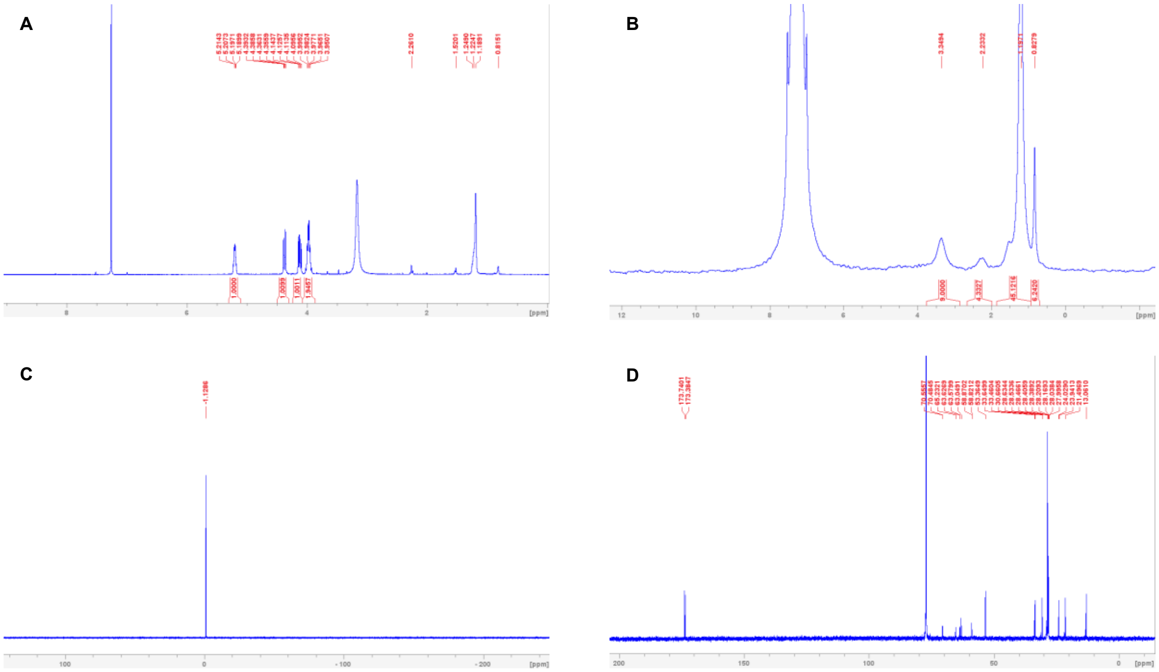


**Figure S7. Spectral analyses of DSPC-$d_{83}$. A.** $^{1}H$ NMR 400 MHz ($CDCl_3$) spectrum. **B.** $^{2}H$ NMR 61.4 MHz ($CDCl_3$) spectrum. **C.** $^{2}H$ NMR 61.4 MHz ($CDCl_3$) spectrum. **D.** $^{13}C$ {$^{1}H$, $^{2}H$} NMR 100.6 MHz spectrum.

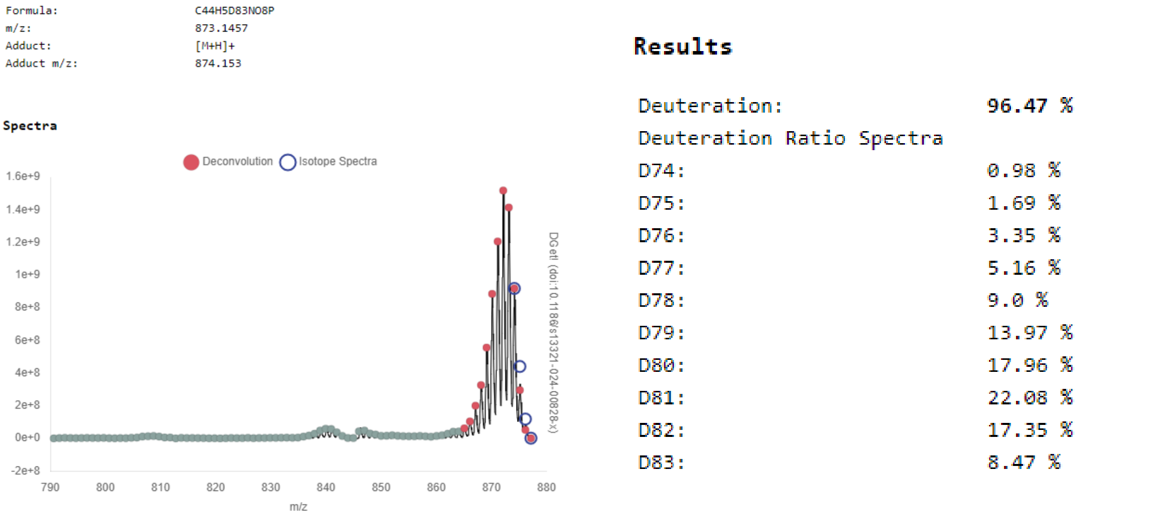


**Figure S8. The use of DGet! For DSPC-$d_{83}$.** DGet! software (https://dget.app/) was employed to calculate the average isotopic purity of DSPC-$d_{83}$ using the distribution of peaks of the $[M+H]^+$ adduct. The isotopic purity for the molecule at the indicated positions (tail and head, $d_{83}$) was quoted as 96.5±2% based on this calculation and an arbitrary error presumption.

4-(Dimethylamino)-butanoic acid, (10Z,13Z)-1-(9Z,12Z)-9,12-octadecadien-1-yl-10,13-nonadecadien-1-yl ester-$d_{62}$ (**DLin-MC3-DMA-$d_{62}$ average 96%D**)

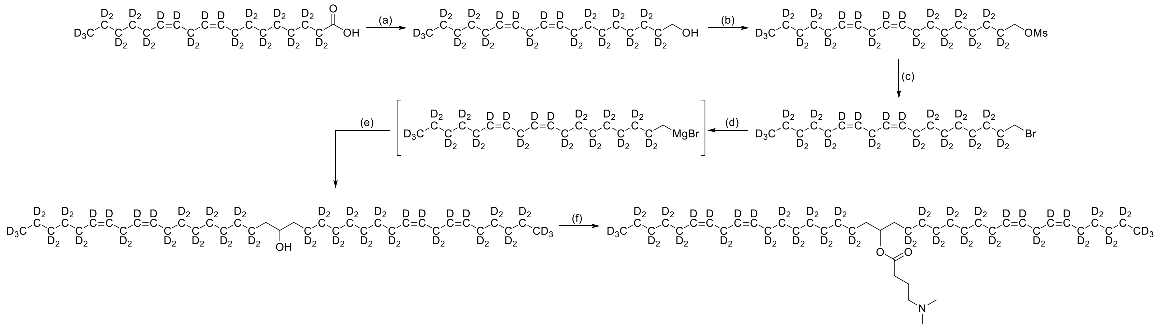

**Scheme S2. Synthesis steps of DLin-MC3-DMA-$d_{62}$.** Reagents and conditions: (a) $LiAlH_4$, THF, 0 °C to room temperature, 16 h; (b) MsCl, $Et_3N$, $CH_2Cl_2$, 0 °C to room temperature, 16 h; (c) $MgBr_2.OEt_2$, $Et_2O$, reflux, 18 h; (d) Mg, 1,2-dibromoethane, $Et_2O$, reflux, 1.5 h; (e) i) Ethyl formate, $Et_2O$, room temperature, 16 h, ii) NaOMe, MeOH, room temperature, 16 h; (f) 4-Dimethylaminobutanoic acid hydrochloride, $iPr_2EtN$, DMAP, EDCI, $CH_2Cl_2$, room temperature, 16 h.

(9*Z*,12*Z*)-Octadeca-9,12-dien-1-ol-$d_{31}$

Lithium aluminium hydride (300 mg, 8.02 mmol) was suspended in anhydrous tetrahydrofuran (10 ml) and cooled to 0 °C. Linoleic acid-$d_{31}$ (1 g, 3.21 mmol) (5) dissolved in tetrahydrofuran (5 ml) was added dropwise and the mixture was warmed to room temperature and stirred for 16 hours. The suspension was cooled to 0 °C and quenched with water (0.3 ml), aqueous sodium hydroxide (15%, 0.3 ml) and further water (0.9 ml). The mixture was stirred for 15 minutes at room temperature, before anhydrous magnesium sulfate was added and the mixture stirred for a further 10 minutes. The suspension was filtered through a pad of Celite and the filtrate concentrated under reduced pressure. The residue was passed through a plug of silica, eluting with ethyl acetate, hexane (0:1 to 3:17) to obtain the title compound (0.95 g, 99%) as a light-yellow oil. $R_f$ = 0.3 (3:17 EtOAc, hexane; Hanessian's); $^1H$ NMR (400 MHz, $CDCl_3$) δ 5.34 (residual), 3.62 (2H, s), 2.0 (residual), 1.57-1.40 (residual), 1.34-1.20 (residual), 0.83 (residual) ppm; $^2H$ NMR (61.4 MHz, $CDCl_3$) δ 5.48-5.29 (4D, m), 2.74 (2D, bs), 2.00 (4D, bs), 1.53 (2D, bs), 1.39-1.14 (16D, m), 0.84 (3D, s) ppm; $^{13}C\{^1H, ^2H\}$ NMR (101 MHz, $CDCl_3$) δ 129.8, 129.7, 127.6, 127.5, 63.1, 31.9, 30.3, 28.6, 28.3, 28.3, 28.2, 28.0, 26.3, 24.9, 24.6, 21.4, 13.0 ppm.

(9*Z*,12*Z*)-Octadeca-9,12-dien-1-yl methanesulfonate-$d_{31}$

The alcohol (940 mg, 3.16 mmol) was dissolved in anhydrous dichloromethane (10 ml) and cooled to 0 °C. Triethylamine (0.57 ml, 4.11 mmol) was added, followed by methanesulfonyl chloride (0.27 ml, 3.47 mmol), dropwise. The mixture was heated to room temperature and stirred for 16 hours. The reaction mixture was diluted with dichloromethane (20 ml), washed with water (30 ml), aqueous saturated sodium hydrogen carbonate (30 ml), dried over sodium sulfate and concentrated under reduced pressure. The crude residue was passed through a plug of silica, eluting with ethyl acetate, hexane (0:1 to 3:17) to obtain the title compound (1.16 g, 97%) as a faint yellow oil. $R_f$ = 0.3 (3:17 EtOAc, hexane; Hanessian's); $^1H$ NMR (400 MHz, $CDCl_3$) δ 5.34 (residual), 4.21 (2H, s), 3.00 (3H, s), 2.0 (residual), 1.72 (residual) 1.36-1.20 (residual), 0.83 (residual) ppm; $^2H$ NMR (61.4 MHz, $CDCl_3$) δ 5.51-5.26 (4D, m), 2.74 (2D, bs), 2.00 (4D, bs), 1.72 (2D, bs), 1.43-1.13 (16D, m), 0.84 (3D, s) ppm; $^{13}C\{^1H, ^2H\}$ NMR (101 MHz, $CDCl_3$) δ 129.8, 129.6, 127.7, 127.5, 70.2, 30.3, 28.5, 28.3, 28.2, 28.1, 28.0, 27.8, 26.3, 26.2, 24.9, 24.3, 21.4, 13.0 ppm.

(6Z,9Z)-18-Bromooctadeca-6,9-diene-$d_{31}$

The mesylate (1.16 g, 3.09 mmol) was dissolved in anhydrous diethyl ether (40 ml) and treated with magnesium bromide diethyl etherate complex (2.39 g, 9.26 mmol). The suspension was heated at reflux for 18 hours, before it was cooled to 0 °C and quenched with ice-water (100 ml). The layers were separated and the organic phase washed with aqueous potassium carbonate solution (1% wt/v, 100 ml), brine (100 ml), dried over sodium sulfate and concentrated under reduced pressure. The crude product was purified by flash column chromatography using hexane as an eluent to obtain the title compound (988 mg, 89%) as a colourless oil. $R_f$ = 0.7 (hexane, Hanessian's); $^1$H NMR (400 MHz, $CDCl_3$) δ 5.35 (residual), 3.39 (2H, s), 2.0 (residual), 1.81 (residual) 1.43-1.20 (residual), 0.83 (residual) ppm; $^2$H NMR (61.4 MHz, $CDCl_3$) δ 5.56-5.23 (4D, m), 2.74 (2D, bs), 2.00 (4D, bs), 1.82 (2D, bs), 1.49-1.08 (16D, m), 0.84 (3D, s) ppm; $^{13}$C{$^1$H, $^2$H} NMR (101 MHz, $CDCl_3$) δ 129.8, 129.6, 127.6, 127.5, 34.0, 32.0, 30.3, 28.5, 28.3, 28.1, 28.0, 27.6, 27.1, 26.3, 26.3, 24.9, 21.4, 13.1 ppm.

(6Z,9Z,28Z,31Z)-Heptatriaconta-6,9,28,31-tetraen-19-ol-$d_{62}$

A dried flask was charged with magnesium turnings (82 mg, 3.36 mmol). The flask was evacuated and backfilled with nitrogen. Anhydrous diethyl ether (0.5 ml) was added followed by 1,2-dibromoethane (1-2 drops). The mixture was heated at reflux for 10 minutes before a solution of the alkyl bromide (0.97 g, 2.69 mmol) in diethyl ether (1 ml) was added dropwise. The mixture was heated at reflux for 1.5 hours. The reaction mixture was cooled to 0 °C, treated with a solution of ethyl formate (0.097 ml, 1.21 mmol) in diethyl ether (1 ml), slowly over 20 minutes, warmed to room temperature and stirred for 16 hours. The mixture was cooled to 0 °C and quenched with water (2 ml), followed by diluted sulfuric acid (1 M, 10 ml). The biphasic solution was transferred to a separating funnel and extracted with diethyl ether (3 x 10 ml). The combined extracts were dried over sodium sulfate and concentrated under reduced pressure. The crude residue was taken up in methanol (10 ml) and treated with sodium methoxide (73 mg, 1.35 mmol). The mixture was stirred at room temperature for 16 h before volatiles were removed under a stream of nitrogen. The residue was diluted with hydrochloric acid (1 M, 15 ml), extracted with diethyl ether (3 x 20 ml), dried over sodium sulfate and concentrated under reduced pressure. The crude product was purified by flash column chromatography using diethyl ether, hexane (0:1 to 1:9) as an eluent to obtain the title compound (0.56 g, 78%) as a faint yellow oil. $R_f$ = 0.3 (1:9 diethyl ether, hexane, Hanessian's); $^1$H NMR (400 MHz, $CDCl_3$) δ 5.35 (residual), 3.61-3.53 (1H, m), 1.50 (1H, bs), 1.49-1.33 (4H, m), 1.32-1.18 (residual), 0.83 (residual) ppm; $^2$H NMR (61.4 MHz, $CDCl_3$) δ 5.57-5.23 (8D, m), 2.74 (4D, bs), 2.01 (8D, bs), 1.47-1.01 (36D, m), 0.84 (6D, s) ppm; $^{13}$C{$^1$H, $^2$H} NMR (101 MHz, $CDCl_3$) δ 129.8, 129.7, 127.6, 127.5, 72.2, 37.4, 30.3, 28.7, 28.6, 28.4, 28.3, 28.3, 28.1, 26.3, 26.3, 24.9, 24.8, 21.4, 13.1 ppm.

D-Lin-MC3-DMA-$d_{62}$

The alcohol (550 mg, 0.93 mmol) was dissolved in anhydrous dichloromethane (3.7 ml) and treated with 4-dimethylaminobutanoic acid hydrochloride (187 mg, 1.12 mmol), N,N-diisopropylethylamine (0.44 ml, 2.51 mmol) and 4-dimethylaminopyridine (17 mg, 0.14 mmol). The mixture was stirred for 5 minutes before N-(3-dimethylaminopropyl)-N′-ethylcarbodiimide hydrochloride (267 mg, 1.4 mmol) was added. The reaction mixture was stirred for 16 hours before it was quenched with saturated aqueous sodium hydrogen carbonate solution (6 ml). The biphasic mixture was transferred to a separatory funnel and extracted with dichloromethane (4 x 20 ml). The combined organic extracts were dried over sodium sulfate and concentrated under reduced pressure. The crude product was purified by flash column chromatography eluting with methanol, chloroform (0:1 to 1:19) to obtain the title compound (490 mg, 75%) as a faint yellow oil. $R_f$ = 0.4 (1:19 methanol, dichloromethane, Hanessian's); $^1$H NMR (400 MHz, $CDCl_3$) δ 5.35 (residual), 4.85 (1H, quin., J =

6.3 Hz), 2.72 (residual), 2.33 (4H, q, J = 6.2 Hz), 2.26 (6H, s), 1.99 (residual), 1.81 (2H, quin., J = 7.3 Hz), 1.48 (4H, d, J = 6.2 Hz), 1.34-1.17 (residual), 0.83 (residual) ppm; $^{2}H$ NMR (61.4 MHz, $CDCl_3$) δ 5.54-5.26 (8D, m), 2.74 (4D, bs), 2.00 (8D, bs), 1.46-1.00 (36D, m), 0.84 (6D, s) ppm; $^{13}C\{^{1}H\}$ NMR (101 MHz, $CDCl_3$) δ 173.4, 74.5, 58.9, 45.4, 34.0, 32.5, 23.0 ppm; $^{13}C\{^{1}H, ^{2}H\}$ NMR (101 MHz, $CDCl_3$) δ 173.4, 129.8, 129.7, 127.6, 127.5, 74.5, 58.9, 45.4, 34.0, 32.5, 30.3, 28.6, 28.4, 28.3, 28.3, 28.3, 28.1, 26.3, 26.3, 24.9, 24.5, 23.0, 21.4, 13.1 ppm; MS (ESI) Calc. $C_{43}H_{17}D_{62}NO_2$ $[M+H]^+$: 705.0081. Deuteration level calculated as 95.8% using DGet!.

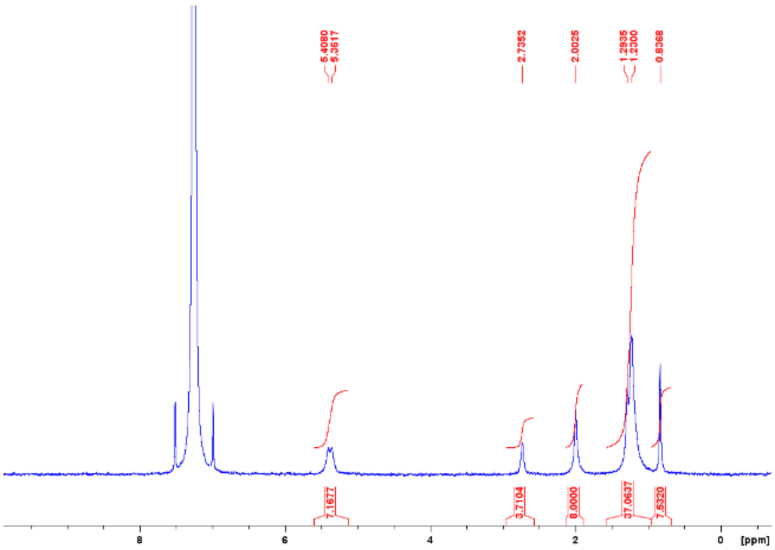


**Figure S9. Spectral analyses of MC3-$d_{62}$.** $^{2}H$ NMR 61.4 MHz ($CDCl_3$) spectrum.

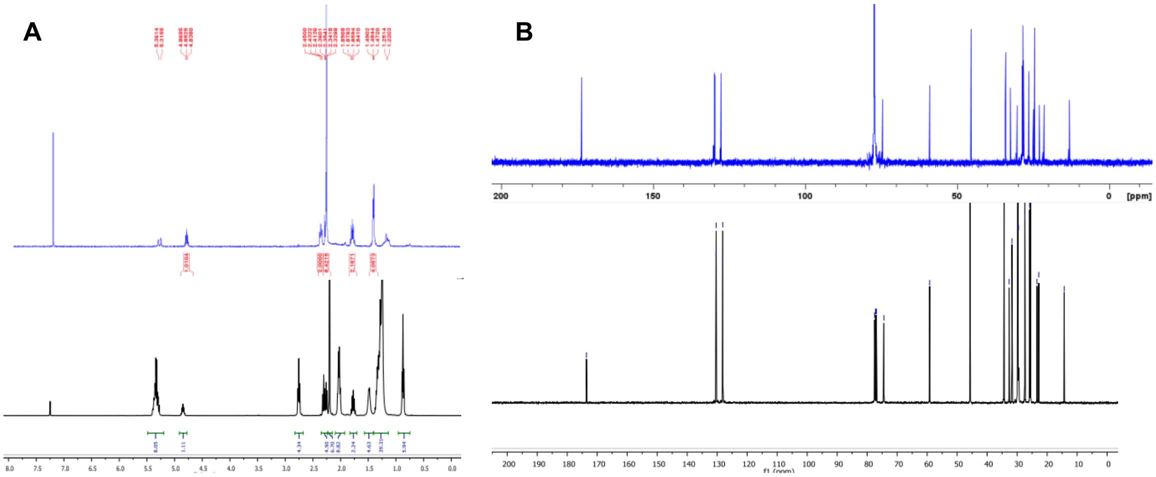


**Figure S10. Spectral analyses of MC3-$d_{62}$ in comparison to the literature. A.** Overlay of $^{1}H$ NMR 400 MHz ($CDCl_3$) spectrum of MC3-$d_{62}$ (top panel, blue) and $^{1}H$ NMR 400 MHz ($CDCl_3$) spectrum of protiated MC3 (bottom panel, black). **B.** Overlay of $^{13}C$ $\{^{1}H, ^{2}H\}$, 100.6 MHz NMR spectrum of MC3-$d_{62}$ (top, blue) with $^{13}C$ $\{^{1}H\}$, 100.6 MHz NMR spectrum of protiated MC3 (bottom, black). The spectra for protiated MC3 were reported by Jayaraman *et al*. (6).

| | |
|---|---|
| Formula: | C43H17D62NO2 |
| m/z: | 704.0002 |
| Adduct: | [M+H]+ |
| Adduct m/z: | 705.0075 |

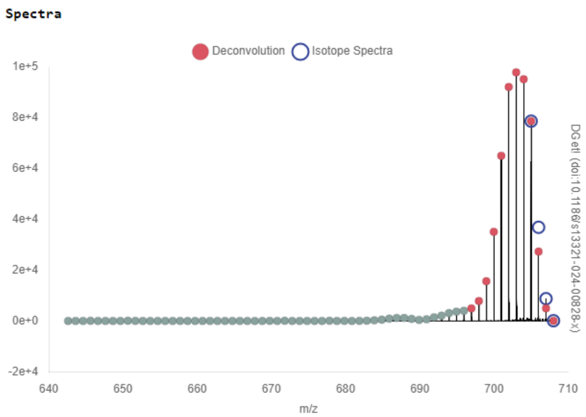


**Results**

| | |
|---|---|
| Deuteration: | 95.78 % |
| Deuteration Ratio Spectra | |
| D54: | 1.04 % |
| D55: | 1.8 % |
| D56: | 3.77 % |
| D57: | 8.64 % |
| D58: | 15.19 % |
| D59: | 19.77 % |
| D60: | 18.63 % |
| D61: | 17.83 % |
| D62: | 13.33 % |

**Figure S11. The use of DGet! MC3-$d_{62}$.** DGet! software (https://dget.app/) was employed to calculate the average isotopic purity of MC3-$d_{62}$ using the distribution of peaks of the $[M+H]^+$ adduct. The isotopic purity for the molecule at the indicated positions (tail, $d_{62}$) was quoted as 95.8±2% based on this calculation and an arbitrary error presumption.

**Cholesterol-$d_{45}$ (average 79%D)**

Synthesised according to previously reported procedures (7). The crude solid was purified by flash chromatography using ethyl acetate, hexane 0:1 to 1:9 as an eluent to provide cholesterol-$d_{45}$ (343 mg) as a white solid, $^1$H NMR (400 MHz, $CDCl_3$) δ 0.64 (m, residual), 0.81–0.90 (m, residual), 0.96–1.00 (m, residual), 1.04–1.11 (m, residual), 1.21–1.34 (m, residual), 1.41–1.51 (m, residual), 1.78–1.84 (m, residual), 1.93–1.99 (m, residual), 2.20–2.30 (m, residual), 3.52 (m, residual), 5.34 (s, residual). $^2$H NMR (61.4 MHz, $CDCl_3$) δ 0.66 (br s), 0.84 (br s), 0.98 (br s), 1.10 (br s), 1.30 (br s), 1.47 (br s), 1.81 (br s), 1.95 (br s), 2.25 (br s), 3.51 (br s), 5.39 (br s). $^{13}$C{$^1$H} NMR (101 MHz, $CDCl_3$) δ 10.8–12.0 (m), 17.1–19.4 (m), 19.7–20.9 (m), 21.4–24.0 (m), 26.5–28.4 (m), 30.6–32.1 (m), 34.7–35.8 (m), 35.9–37.1 (m), 37.9–39.6 (m), 41.5–42.4 (m), 49.0–50.2 (m), 55.2–55.7 (m), 55.7–56.3 (m), 56.5–56.8 (m), 56.9 (s), 71.1–71.9 (m), 121.2–121.9 (m), 140.8–141.1 (m). $^{13}$C{$^1$H, $^2$H} NMR (101 MHz, $CDCl_3$) δ 11.1 (m), 11.4 (m), 11.7 (m), 17.8 (m), 18.2 (m), 18.6 (m), 18.9 (m), 19.2 (m), 20.3 (s), 20.4 (s), 20.6 (m), 20.7 (m), 21.6 (m), 21.8 (m), 22.1 (m), 22.8 (m), 22.9 (m), 23.4 (s), 23.5 (m), 27.3 (m), 27.5 (m), 27.7 (m), 30.8 (m), 31.0 (m), 31.2 (m), 31.7 (m), 34.8 (m), 35.1 (m), 35.5 (m), 36.2 (m), 36.4 (m), 36.8 (m), 37.1 (m), 38.4 (m), 38.8 (m), 39.2 (m), 41.9 (m), 42.3 (m), 49.3 (m), 55.4 (m), 56.0 (m), 56.6 (m), 71.3 (m), 121.4 (m), 141.0 (m). MS (ESI) Calc. $C_{27}HD_{45}O$ $[M-H_2O+H]^+$ Deuteration level calculated as 78.7% using DGet! (3).

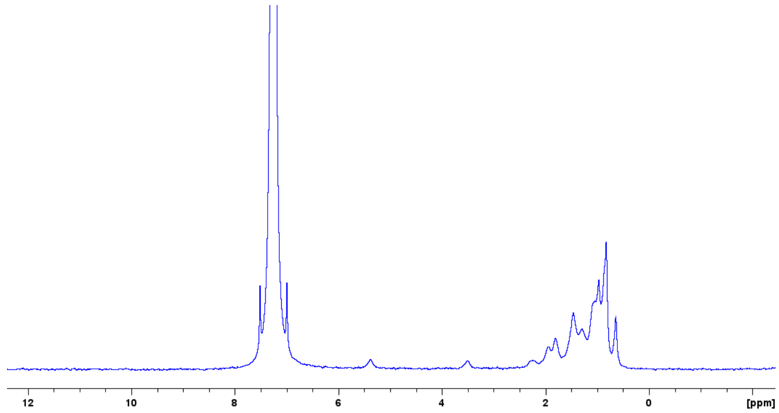


**Figure S12. Spectral analyses of Cholesterol-d$_{45}$.** $^{2}$H NMR 61.4 MHz ($CDCl_3$) spectrum.

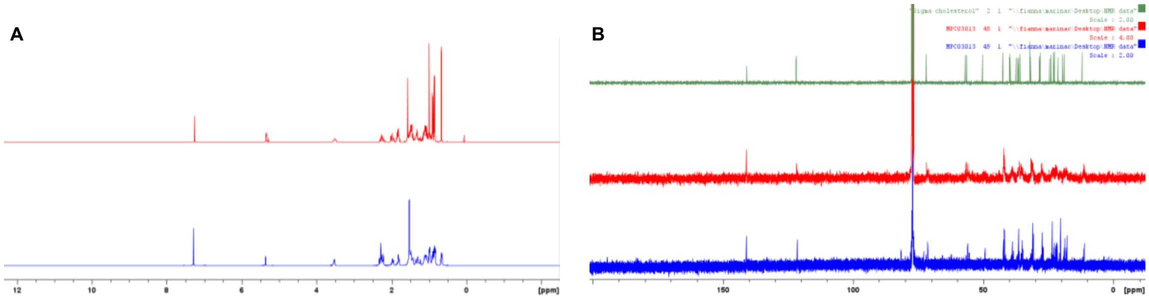


**Figure S13. Spectral comparison of cholesterol-d$_{45}$. A.** Stacked view of $^{1}$H NMR (400 MHz, $CDCl_3$) spectrum of protiated cholesterol (top panel, red) and deuterated cholesterol-d$_{45}$ (bottom panel, blue). **B.** Stacked view of $^{13}$C{$^{1}$H} NMR (101 MHz, $CDCl_3$) spectrum of protiated cholesterol (top panel, green), cholesterol-d$_{45}$ (middle panel, red) and $^{13}$C{$^{1}$H, $^{2}$H} NMR (101 MHz, $CDCl_3$) spectrum of cholesterol-d$_{45}$ (bottom panel, blue).

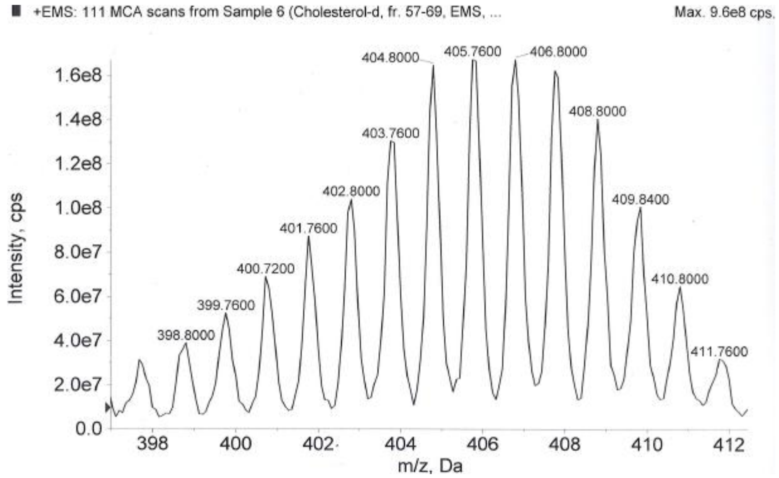


**Figure S14.** ESI mass spectrum of cholesterol-d$_{45}$ showing distribution of isotopologues.

**SANS computations**

**Table S1. Neutron scattering length densities (SLDs) used in SANS fitting.**

| Compound | SLDx10$^{-6}$ (A°$^{-2}$) hydrogenous form | SLDx10$^{-6}$ (A°$^{-2}$) in $D_2O$ deuterated form | Molecular volume (A°$^{3}$) |
|---|---|---|---|
| $H_2O$ | -0.56 | 6.39 | 30 |
| MC3 | 0.09 | 5.03 | 1290 |
| Cholesterol | 0.21 | 5.77 | 630 |
| DSPC | 0.18 | 6.52 | 1322 |
| DMG-$PEG_{2k}$ | 0.12 | - | 970 (only DMG) |
| RNA | 3.6 | - | 325 |

SasView was used as the primary analysis software. To capture the varying amount of solvent in the core or shell, we used a custom core-shell model 'core_shell_fuzzy_sphere'. Core_shell_fuzzy_sphere model parametrizes the original SLD variables from the built-in core-shell model into the two fitting variables: i. volume fraction of solvent ( $v_{sol}$ ) and ii. SDL of the core comprising of only lipids in the core ( $SLD_{dry\ core}$ ) or SDL of the shell comprising of only lipids in the shell ( $SLD_{dry\ shell}$ ), excluding solvent. Therefore, we compute $SLD_{core} = v_{sol}SLD_{sol} + \left(1 - v_{sol}\right)SLD_{dry\ core}$ and $SLD_{shell} = v_{sol}SLD_{sol} + \left(1 - v_{sol}\right)SLD_{dry\ shell}$ for the SDL of the core and SDL of the shell, respectively, where $SLD_{sol}$ is the scattering length density of the solvent. $SLD_{dry\ core}$ and $SLD_{dry\ shell}$ are weighted averages of the SLDs for all lipids present in the corresponding core and shell.

We first performed a simultaneous fit on 2-5 curves using core_shell_fuzzy_sphere model with constant values for the scale factor, background, and $SLD_{sol}$ . The scale factor (set at 0.003 in this work) was determined by calculating the volume fraction of the LNPs, derived from the final lipid concentration and lipid molecular volumes. The background was subtracted during data processing and set to a minimal margin of error. $SLD_{sol}$ was determined with the respective $H_2O/D_2O$ ratio. Given 5 vol.% ethanol in the final LNP solutions, $SLD_{sol}$ is computed for different $D_2O$ volume factions as 6.05x10$^{-6}$, 3.93x10$^{-6}$, 2.74x10$^{-6}$, 1.82x10$^{-6}$, and 0.63x10$^{-6}$ A°$^{-2}$ for 95, 64.6, 47.5, 34.2, and 17.1 vol.% $D_2O$.

The core radius, shell thickness, $SLD_{core}$ and $SLD_{shell}$ were optimized by minimizing the Chi squared for the fitting error, and the radius and thickness were constrained to be equal among different solvent contrasts. To do so, we first hypothesized that solvent could be present in both the core and shell. Since $SLD_{sol}$ varies with $D_2O$ volume faction, we allowed $SLD_{core}$ and $SLD_{shell}$ to vary to account for the potential presence of solvent. Once we find that the fitting error could be minimized by ruling out the presence of solvent, we constrained $SLD_{shell}$ . We therefore constrained $SLD_{shell}$ for all the LNPs but the empty LNPs. To capture the broad peaks around 0.2 A°$^{-1}$, the parameters from the custom core_shell model were transferred to a model containing the sum of the built-in core-shell sphere and broad peak models. Then, each contrast was fit individually, keeping all fitting variable but the breadth and intensity of the broad peak constant.
